# Drosophila Ca_V_2 channels harboring human migraine mutations cause synapse hyperexcitability that can be suppressed by inhibition of a Ca^2+^ store release pathway: Drosophila models of Ca_V_2 migraine mutations

**DOI:** 10.1101/141366

**Authors:** Douglas J. Brusich, Ashlyn M. Spring, Thomas D. James, Catherine J. Yeates, Timothy H. Helms, C. Andrew Frank

## Abstract

Gain-of-function mutations in the human Ca_V_2.1 gene *CACNA1A* cause familial hemiplegic migraine type 1 (FHM1). To characterize cellular problems potentially triggered by Ca_V_2.1 gains of function, we engineered mutations encoding FHM1 amino-acid substitutions S218L (SL) and R192Q (RQ) into transgenes of *Drosophila melanogaster* Ca_V_2/*cacophony*. We expressed the transgenes pan-neuronally. Phenotypes were mild for RQ-expressing animals. By contrast, single mutant SL- and complex allele RQ,SL-expressing animals showed overt phenotypes, including sharply decreased viability. By electrophysiology, SL- and RQ,SL-expressing neuromuscular junctions (NMJs) exhibited enhanced evoked discharges, supernumerary discharges, and an increase in the amplitudes and frequencies of spontaneous events. Some spontaneous events were gigantic (10-40 mV), multi-quantal events. Gigantic spontaneous events were eliminated by application of TTX – or by lowered or chelated Ca^2+^ – suggesting that gigantic events were elicited by spontaneous nerve firing. A follow-up genetic approach revealed that some neuronal hyperexcitability phenotypes were reversed after knockdown or mutation of Drosophila homologs of phospholipase Cβ (PLCβ), IP_3_ receptor, or ryanodine receptor (RyR) – all factors known to mediate Ca^2+^ release from intracellular stores. Pharmacological inhibitors of intracellular Ca^2+^ store release produced similar effects. Interestingly, however, the decreased viability phenotype was not reversed by genetic impairment of intracellular Ca^2+^ release factors. On a cellular level, our data suggest inhibition of signaling that triggers intracellular Ca^2+^ release could counteract hyperexcitability induced by gains of Ca_V_2.1 function.

**AUTHOR SUMMARY:** Prior research has demonstrated that gain-of-function mutations in a gene important for neurotransmission (*CACNA1A*) are known to cause migraine in humans. We attempted to mimic some of those gain-of-function mutations in a simple genetic model organism and to examine neurotransmission by electrophysiology. Our findings yield potential clues as to how particular migraine-causing mutations may impact neurophysiology on a cellular level. We used the fruit fly *Drosophila melanogaster* and its model synapse, the neuromuscular junction (NMJ) to perform our studies. We document three main advances: 1) characterization of fruit fly models harboring gain-of-function calcium channel alterations known to cause human familial hemiplegic migraine type 1 (FHM1); 2) characterization of hyperactive neurotransmission caused by one of these alterations; and 3) an ability to quell hyperactive neurotransmission by impairing intracellular Ca^2+^ store release, through both genetic and pharmacological means. Our work contributes to a broader understanding of how pathological mutations could impact cellular physiology. More generally, the utilization of genetic model organisms promises to uncover potential ways to reverse those impacts.

## INTRODUCTION

Episodic neurological disorders like migraine, epilepsy, and ataxia can result from underlying ion channel dysfunctions (1–3). For many such disorders, little is known about how aberrant channel functions affect neuronal signaling paradigms. Cell-based and model organism-based examinations of disease-causing mutations could offer insights into disease-relevant biological processes. One Mendelian form of migraine – familial hemiplegic migraine type 1 (FHM1) – results from gain-of-function missense mutations in human *CACNA1A*, which encodes the α1 subunit of Ca_V_2.1 (P/Q)-type calcium channels (4). Two FHM1-causing amino-acid substitutions alter highly conserved Ca_V_2.1 α1 amino-acid residues, R192 and S218 (4, 5). The R192Q amino-acid substitution (RQ) causes “pure” FHM1, while the S218L substitution (SL) causes a severe combination of FHM1, seizures, and susceptibility to edema following head injury (4, 5). These two FHM1-causing amino-acid substitutions have been studied intensely (6), most notably in knock-in mouse models of FHM1 (7–9).

FHM1 knock-in mice display gain-of-function Ca_V_2.1 phenotypes at neurons and synapses. Model synapses studied include the diaphragm neuromuscular junction (NMJ) (10, 11), the calyx of Held (12–14), the trigeminal sensory neuron pathway (15–17), and cortical neurons (18, 19). At the mouse NMJ, both RQ and SL increase the frequency of spontaneous excitatory potentials (10, 11). These increases in quantal frequency are dependent on mutation dose and are more pronounced in SL versus RQ. SL also elicits broadening of evoked end-plate potentials at the mouse NMJ (11). At the calyx of Held, both substitutions result in enhanced excitatory postsynaptic currents (EPSCs) (12–14), and it has been reported that SL causes an increase in the resting intracellular neuronal calcium, which could be responsible for some potentiation of synapse function (12).

It was recently reported that 2,5′-di(tertbutyl)-1,4,-benzohydroquinone (BHQ) reverses aspects of SL-induced gating dysfunction and short-term plasticity (20). As part of that study, we found that BHQ also restores short-term synaptic plasticity to NMJs in fruit fly larvae expressing a transgene that encodes an S161L amino-acid substitution in Drosophila Ca_V_2/Cacophony – the functional equivalent of human Ca_V_2.1 S218L (20). Independent follow-up work in the mouse S218L model demonstrated that BHQ application also blunts cortical spreading depression susceptibility (21). Given these collective results, a further examination of fruit fly synapses could be valuable for uncovering relevant molecular and electrophysiological consequences of Ca_V_2.1 gains of function.

For the present study, we characterized the fruit fly as a way to model neuronal effects of FHM1-causing mutations. We neuronally expressed *cacophony* transgenes harboring the *Drosophila melanogaster* equivalents of RQ or SL – or both RQ and SL concurrently (denoted as “RQ,SL”). On the organismal level, neuronal expression of SL or RQ,SL transgenes drastically impaired overall health. On the synapse level, SL and RQ,SL transgenes markedly enhanced aspects of evoked and spontaneous neurotransmission, consistent with prior studies in mice. Through a combination of genetics, RNA interference, pharmacology, and electrophysiology, we uncovered evidence that impairment of a conserved intracellular signaling pathway that triggers store Ca^2+^ release reverses hyperexcitability phenotypes in the context of gain-of-function Drosophila Ca_V_2.

## RESULTS

### Transgenic Drosophila Ca_V_2 “FHM1” channels cause coarse larval phenotypes and fly lethality

We utilized *Drosophila melanogaster* to study the impact that FHM1-inducing Ca_V_2.1 amino-acid substitutions may exert on the level of individual synapses. Drosophila *cacophony* encodes the α1 subunit of fruit fly Ca_V_2-type channels. We cloned two amino-acid substitutions that cause human FHM1 (S218L and R192Q) into the analogous codons of a functional Drosophila *UAS-cacophony (cac)-eGFP* transgene (22). Single mutant transgenes were termed “SL” (*UAS-cac-eGFP^S161L^*) (20) or “RQ” (*UAS-cac-eGFP^R135Q^*) (Fig. 1A). We also generated a transgene containing both mutations in *cis* on the same cDNA clone, termed “RQ,SL” (*UAS-cac-eGFP^R135Q, S161L^*). This is not a naturally occurring allele in humans with FHM1. We reasoned *a priori* that this complex allele could yield a genetically sensitized background for Ca_V_2 gain-of-function in Drosophila. “WT” signifies previously characterized wild-type *UAS-cac-eGFP^WT^* transgenes (22).

**Figure 1:**
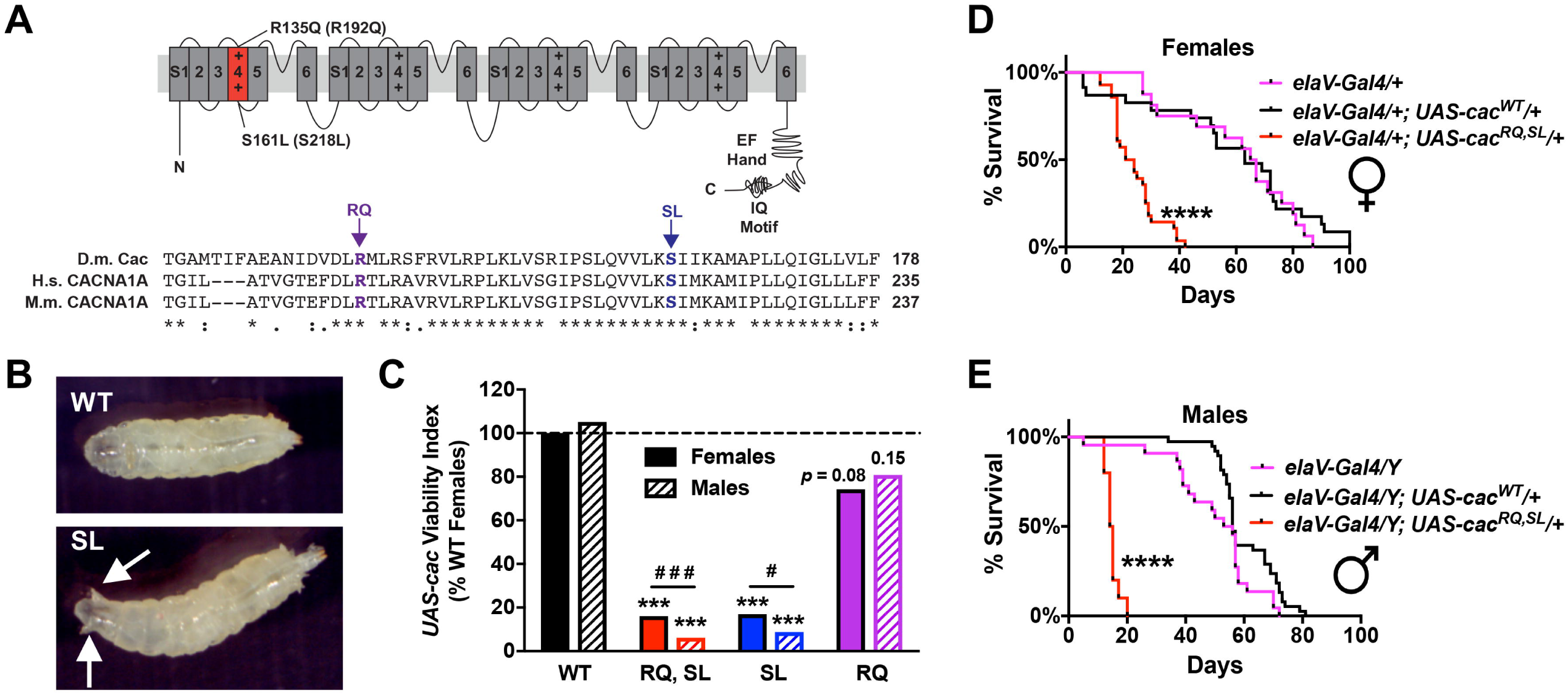
SL- and RQ,SL-expressing flies exhibit coarse phenotypes. **(A)** Schematic of Ca_V_2-type calcium channel α1a subunit, with substitutions to Drosophila Cacophony (Cac) residues indicated (mammalian residues in parentheses) and a CLUSTAL-Omega alignment of Cac, human CACNA1A, and mouse CACNA1A amino acids spanning the relevant region ([*] - fully conserved; [:] - strongly similar; [.] -weakly similar). **(B, C)** Visible phenotypes for larvae resulting from crosses of *elaV(C155)-Gal4* females x *Balancer*/*UAS-cac-eGFP^MUT or WT^* males. **(B)** Premature spiracle protrusion in a larva expressing the *UAS-cac-eGFP^SL^* transgenic line (also observed with *UAS-cac-eGFP^RQ,SL^* expression). The spiracle phenotype did not occur in larvae expressing *UAS-cac-eGFP^RQ^* or *UAS-cac-eGFP^WT^*. **(C)** Same crosses as in (B) showing diminished *UAS-cac-eGFP* mutant viability. “*UAS-cac* Viability Index” = # *UAS-cac-eGFP* transgenic adult progeny/# Balancer Chromosome siblings, normalized to 100% for WT female progeny counts (Table 1 for raw counts; for all comparisons, *n* ≥ 115 Balancer sibling progeny were counted). *** *p* < 0.001 by Fisher’s Exact test compared to WT sex-specific control. # *p* = 0.05, ### *p* < 0.001 by Fisher’s Exact test between sexes for the SL or RQ,SL genotypes. **(D, E)** For both females (D) and males (E), there was starkly diminished longevity for adult flies expressing the RQ,SL transgene. **** *p* < 0.0001 by Log-rank test.

We expressed WT, RQ, SL, and RQ,SL *UAS-cac-eGFP* transgenes in post-mitotic Drosophila neurons using the *elaV(C155)-Gal4* driver and the *Gal4/UAS* expression system (23, 24). We examined transgenic animals qualitatively for visible phenotypes. Neuronal expression of either SL or RQ,SL caused larvae to move in a jerky, uncoordinated manner. At the early third instar stage, SL- and RQ,SL-expressing animals developed protruding, anterior spiracles prematurely – well before the normal time point of wandering third instar stage and pupation (Fig 1B).

Our initial observations indicated that SL- and RQ,SL-expressing animals were not present in expected Mendelian proportions. For each transgene (WT, RQ, SL, and RQ,SL), we set up test crosses (*elaV(C155)-Gal4* females x *Balancer Chromosome*/*UAS-cac-eGFP* males) and counted the number of transgenic *UAS-cac-GFP*-expressing adult progeny and the number of sibling flies carrying a balancer chromosome. We also set up *Gal4* and balancer chromosome control crosses lacking any *UAS-cac-eGFP* transgenes (Table 1). Compared to animals expressing the WT transgene, viability was dramatically diminished for animals expressing the SL and RQ,SL transgenes (Fig. 1C, Table 1). It was also diminished for SL- and RQ,SL-expressing animals compared to genetically matched control siblings carrying the *elaV(C155)-Gal4* driver and a balancer chromosome (Table 1). WT- and RQ-expressing animals did not show significant defects in viability or statistical differences from the *Gal4* control cross (Fig. 1C, Table 1). As expected, there was some depressed viability in animals carrying a balancer chromosome alone (Table 1).

Sex or dose of the SL and RQ,SL transgenes could influence viability. In Drosophila, X-linked dosage compensation equalizes the expression of X-linked genes by doubling X-linked gene transcription in males (25–27). The X-linked neuronal enhancer trap *Gal4* line *elaV(C155)-Gal4* should be expressed at higher levels in hemizygous *elaV(C155)-Gal4/Y* males than in heterozygous *elaV(C155)-Gal4*/+ females. Thus, effects of driving *UAS* transgenes could be stronger in males. Counting male vs. female progeny of SL- and RQ,SL-expressing flies revealed that while viability was starkly diminished for both sexes, it was also significantly lower in SL- and RQ,SL-expressing males than in SL- and RQ,SL-expressing females (Fig. 1C, Table 1).

**Table 1.** Test Crosses and Survival of Adult Progeny. Crosses were performed utilizing *elaV(C155)-Gal4* virgin females x *w/Y; Balancer/* “*(UAS-cac-eGFP or +)*” males. Male and female progeny were counted separately. As expected for fruit fly balancer chromosomes, there was some lethality associated with inheriting a balancer chromosome. Separately, there was profound lethality associated with inheriting either the *UAS-cac-eGFP^RQ,SL^* or *UAS-cac-eGFP^SL^* transgenes driven by pan-neuronal *elaV(C155)-Gal4*. Balancers used depended on the cross and transgene being balanced (*CyO-GFP* was used for “+” and for “*UAS-cac-eGFP^RQ,SL^*” and TM6b was used for the others). Normalized Viability Index scores were calculated from the non-Balancer/Balancer progeny ratio of a single sex; this value was then normalized against the female progeny ratio for “*UAS-cac-eGFP^WT^*” – i.e., 190/115 was set as normalized baseline value = 100.0. For statistical analyses, raw progeny counts were used. *** *p* < 0.001 by Fisher’s exact test, compared to sex-matched progeny counts, utilizing the *UAS-cac-eGFP^WT^* as the control. ^###^ *p* < 0.001, ^#^ *p* = 0.05 by Fisher’s exact test comparing progeny counts of SL- and RQ,SL- expressing males vs. SL- and RQ,SL-expressing females respectively.

| *elaV(C155)-Gal4*<br>x<br>*Balancer / “ ”* | Count<br>(Mendelian<br>exp. per<br>category) | Female Progeny |  | Male Progeny |  | Normalized<br>Viability<br>Index for “ ”<br>(female) | Normalized<br>Viability<br>Index for “ ”<br>(male) |
| --- | --- | --- | --- | --- | --- | --- | --- |
|  |  | *Gal4/+;*<br>“ ”/+ | *Gal4/+;*<br>*Balancer/+* | *Gal4/Y;*<br>“ ”/+ | *Gal4/Y;*<br>*Balancer/+* |  |  |
| “+” | 170 (42.5) | 52 (30.6%) | 37 (21.8%) | 53 (31.2%) | 28 (16.5%) | 85.1 | 114.6 |
| “*UAS-cac-eGFP<sup>WT</sup>*” | 628 (157) | 190 (30.3%) | 115 (18.3%) | 205 (32.6%) | 118 (18.8%) | 100.0 | 105.2 |
| “*UAS-cac-eGFP<sup>RQ,SL</sup>*” | 1040 (260) | 132 (12.7%) | 499 (48.0%) | 38 (3.7%) | 371 (35.7%) | 16.0 *** | 6.2 *** ### |
| “*UAS-cac-eGFP<sup>SL</sup>*” | 308 (77) | 38 (12.3%) | 136 (44.2%) | 17 (5.5%) | 117 (38.0%) | 16.9 *** | 8.8 *** # |
| “*UAS-cac-eGFP<sup>RQ</sup>*” | 542 (135.5) | 157 (29.0%) | 128 (23.6%) | 147 (27.1%) | 110 (20.3%) | 74.2 | 80.9 |

We also assessed adult fly longevity, comparing WT and RQ,SL transgenic flies (Figs. 1D, E). For females, transgenic WT (mean survival: 63 days, *n* = 23) and driver control *elaV(C155)-Gal4/+* animals (66 days, *n* = 16) did not differ with respect to survival. Transgenic RQ,SL females (22.5 days, *n* = 28) had severely stunted longevity (Fig. 1D). The results for males were consistent: longevity of transgenic WT males (median survival: 56 days, *n* = 38) and driver control *elaV(C155)-Gal4/Y* animals (54.5 days, *n* = 22) did not differ statistically. By contrast, the survival of transgenic RQ,SL males (14.5 days, *n* = 10) was markedly diminished (Fig. 1E).

### Cac-GFP localizes normally and levels are comparable across transgenic constructs

We investigated why SL- and RQ,SL-expressing animals were showing overt phenotypes. We considered the possibility that excessive quantities of α1 protein generated via the *GAL4/UAS* expression system could reduce viability. Opposing this idea, neuronal overexpression of WT *UAS-cac* transgenes renders no reported structural, behavioral, or electrophysiological abnormalities (22, 28). Moreover, overexpressed Cac-GFP protein efficiently localizes to active zone structures at synapses like the larval neuromuscular junction (NMJ) (20, 22, 29–32).

Using wandering third instar larvae and *elaV(C155)-Gal4* driver, we first checked Cac-GFP localization of several transgenic lines: WT (published line, *UAS-cac-eGFP^786c^*) (22), RQ,SL (*UAS-cac-eGFP^RQ,SL(2M)^*) (this study), SL (*UAS-cac-eGFP^S/L(3-2M)^*) (20), and RQ (*UAS-cac-eGFP^R/Q(1M)^*) (this study). We used an anti-GFP antibody to detect Cac-GFP and co-stained with a monoclonal antibody against the presynaptic ELKS/CAST active zone protein Bruchpilot (Brp) (33). In all cases, Cac-GFP localized as expected in the larval central nervous system (Figs. 2A-D, red channel). It also predominantly localized to presynaptic active zone sites at neuromuscular junction (NMJ), as expected (Figs. 2E-H), consistent with the reports for the original WT constructs (22, 29).

**Figure 2:**
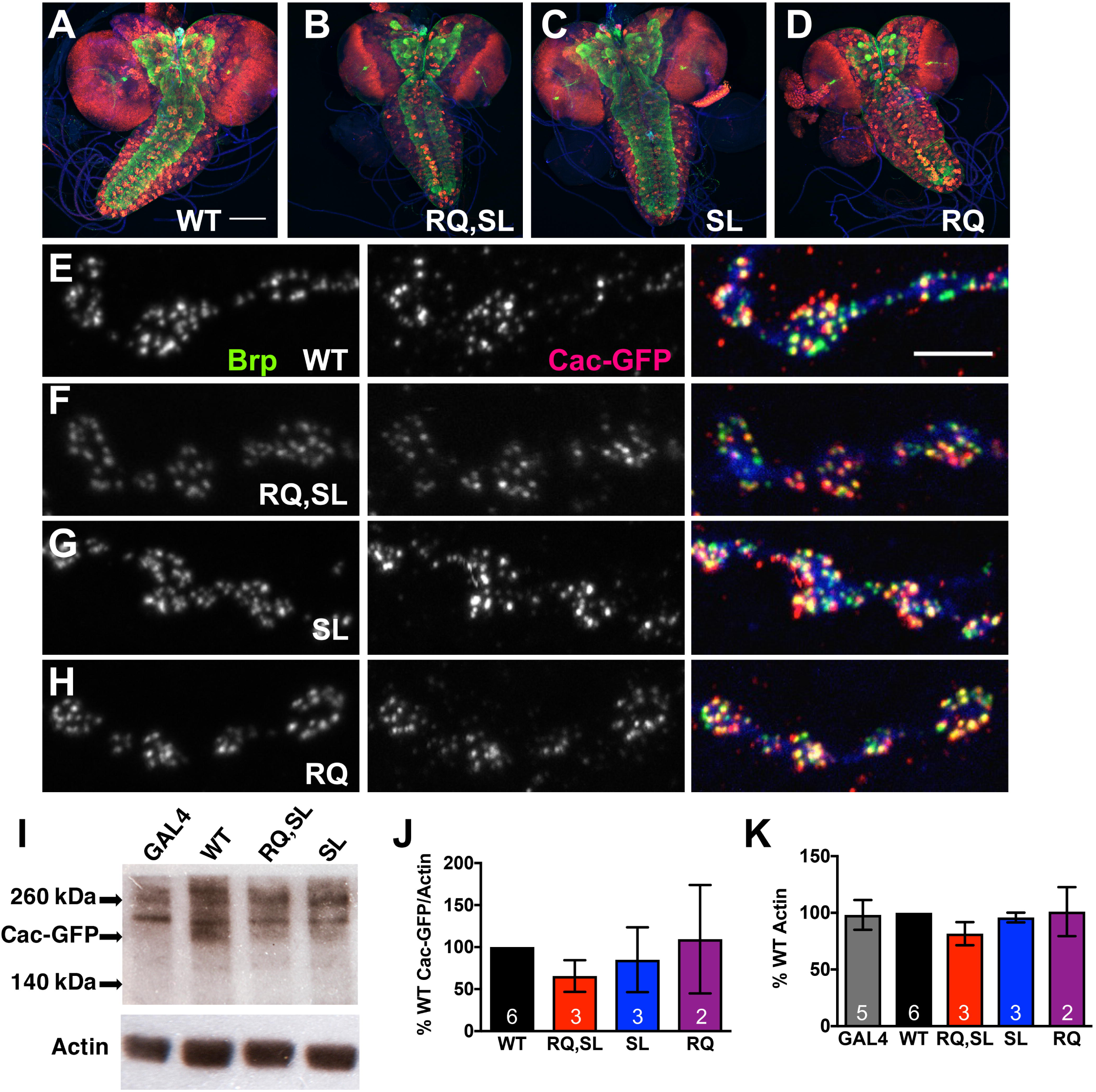
Localization and expression levels of Cac-GFP transgenes are normal. **(A-D)** Images of larval central nervous systems from animals expressing Cac-GFP protein (WT or mutant). Anti-GFP (red), and anti-Bruchpilot (Brp - green) staining are shown. Scale bar 100 μm. **(E-H)** Wild-type and mutant Cac-GFP successfully localized to NMJ active zones, as indicated by co-staining with anti-Brp (green) and anti-GFP (red). Scale bar 5 μm. **(I)** Western blots of fruit fly head lysates (10 heads/lane, single sex per Western), from flies expressing either *elaV-Gal4* alone or the indicated *UAS-cac-eGFP* transgene driven by *elaV-Gal4*. Blots were probed with anti-GFP (top) and anti-Actin (bottom) antibodies. The band corresponding to Cac-GFP is indicated. Other bands are non-specific. **(J)** Compared to WT, there was no statistically significant change in Cac-GFP expression for any of the transgenic lines utilized in this study (band normalized to actin; *p* > 0.65, one-way ANOVA with Dunnett’s multiple comparisons vs. WT; GAL4 alone control excluded from analysis). **(K)** Actin levels were also steady across all transgenic lines (*p* > 0.71, one-way ANOVA with Dunnett’s multiple comparisons vs. WT).

We checked Cac-GFP levels for the different transgenic constructs by Western Blot. We drove the transgenes neuronally using *elaV(C155)-Gal4* and collected adult heads for analysis, blotting for Cac-GFP (239 kDa) with anti-GFP and anti-Actin as a loading control. Compared to *elaV(C155)-Gal4* line controls, each *UAS-cac-eGFP* transgenic line showed an additional, faint band that migrated at a size consistent with Cac-GFP (Fig. 2I). This band was expressed at comparable levels between the WT, RQ, SL, and RQ,SL lines (Fig. 2J), with no appreciable difference in control levels of actin between the lines (Fig. 2K).

### RQ,SL-expressing NMJs show small changes in bouton number and glutamate receptor coverage

Even in the absence of localization or expression-level differences, transgenic mutant Cac-GFP expression could affect synapse growth or development. Previously, we found no significant abnormalities in NMJ synaptic growth for SL-expressing flies (20). We extended our analysis to the RQ,SL transgene line by co-staining third instar larval NMJs with antibodies against the Drosophila PSD-95 homolog, Discs Large (Dlg) and the GluRIIA glutamate receptor subunit (Figs. 3A-D). We observed a very small decrease in the number of Dlg-positive synaptic boutons at RQ,SL-expressing NMJs compared to control WT-expressing NMJs. This decrease was statistically significant only for segment A2, muscle 6/7 (Fig. 3E). We found no significant change in the number of glutamate receptor clusters per NMJ comparing WT-expressing synapses and RQ,SL-expressing synapses (Fig. 3F).

**Figure 3:**
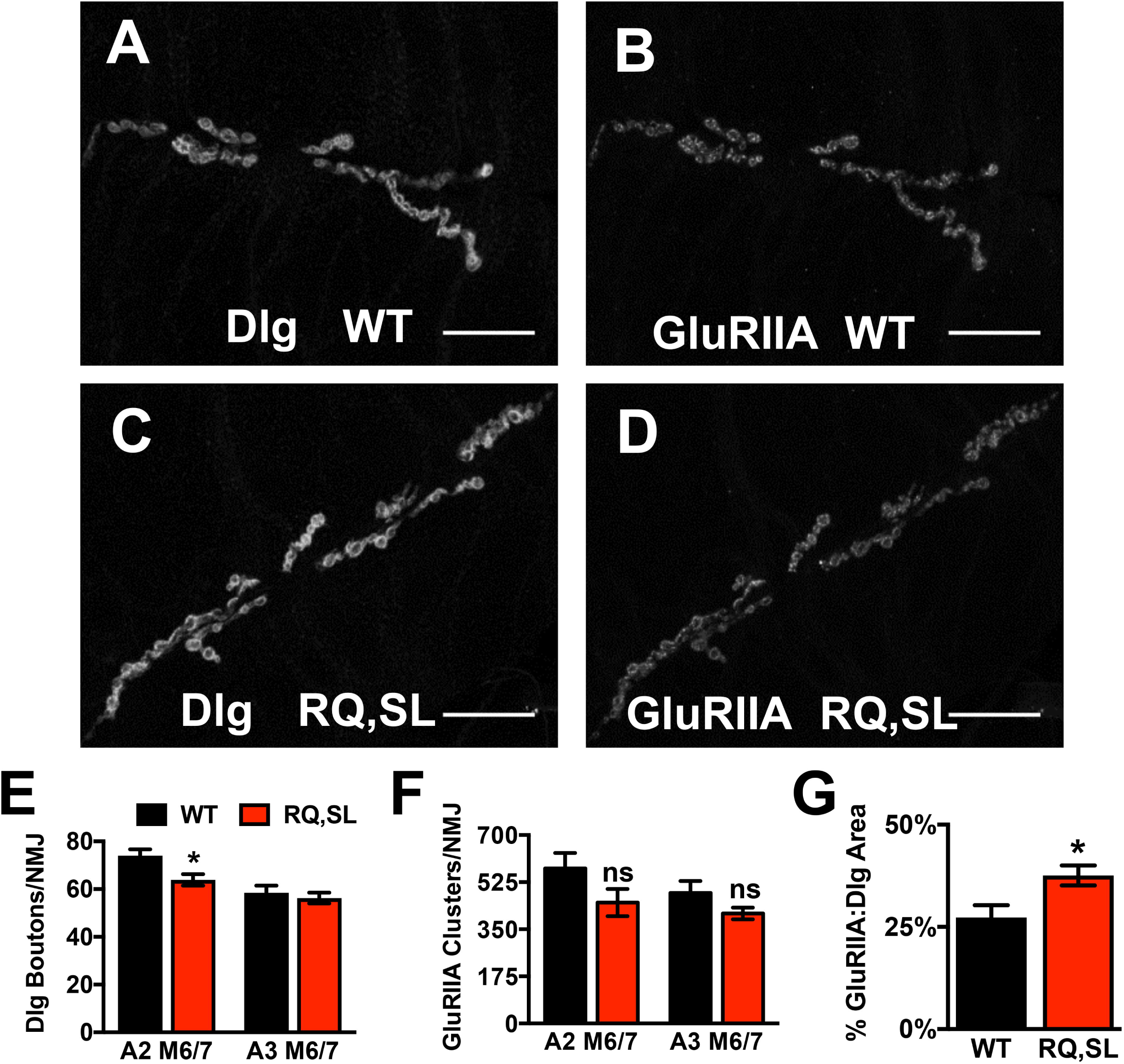
Hallmarks of NMJ development are normal when Cac-GFP transgenes are expressed. **(A-D)** NMJ images of the synapses on Muscle 6/7 of WT- and RQ,SL-expressing third-instar larvae, immunostained with anti-Discs Large (Dlg) and anti-GluRIIA antibodies. Scale bars, 25 μm. **(E)** For RQ,SL-expressing NMJs, average synaptic bouton numbers were normal, except for a slight undergrowth detected for synapse A2 muscle 6/7 (* *p* < 0.05, Student’s T-test vs. WT, *n* ≥ 8 NMJs for all genotypes and segments). **(F)** The number of glutamate receptor clusters per synapse at RQ,SL-expressing NMJs was not statistically significantly different than WT-expressing NMJs (*p* > 0.1, Student’s T-test, *n* ≥ 8 NMJs for all genotypes and segments). **(G)** For RQ,SL-expressing NMJs, there was a small increase in GluRIIA-containing receptor area coverage. (* *p* < 0.05 by Student’s T-test vs. WT for both measures, *n* ≥ 15 NMJs for each genotype).

At RQ,SL-expressing NMJs, the percentage of the synaptic area covered by the GluRIIA clusters – normalized to total Dlg area – was slightly but significantly increased (Fig. 3G). In principle, an expansion of the synaptic area capable of receiving neurotransmitter could underlie gains in synaptic transmission (34). The magnitude of any such change based on this postsynaptic staining profile alone would likely be small but was uncertain based on these measures. We needed to conduct finer analyses by electrophysiology, both to document possible changes in synaptic function and also to test for potential presynaptic contributions when mutant *cac* transgenes were expressed.

### RQ-, SL-, and RQ,SL-expressing NMJs display hyperexcitable evoked synaptic discharges

Coarse phenotypes from neuronally expressed RQ,SL and SL transgenes (Fig. 1) suggested abnormal neuronal or synapse function. Neuronal expression of gain-of-function *UAS-cac-GFP* transgenes could result in enhanced evoked NMJ neurotransmission in Drosophila, similar to the knock-in mouse FHM1 models. Expression of both SL and RQ,SL significantly increased EPSP amplitudes across a range of low extracellular [Ca^2+^] (0.2–0.5 mM) (Fig. 4A, data for 0.4 mM [Ca^2+^]_e_ are shown) (20). Expression of RQ numerically increased average NMJ EPSP amplitudes, but this increase was not statistically significant (Fig. 4A). Neither estimated quantal content (QC) (Fig. 4B) nor calcium cooperativity of release for mutant lines were significantly different than WT across this range of 0.2–0.5 mM [Ca^2+^] (Fig. 4C) (20) (but see more detailed quantal analyses later).

**Figure 4:**
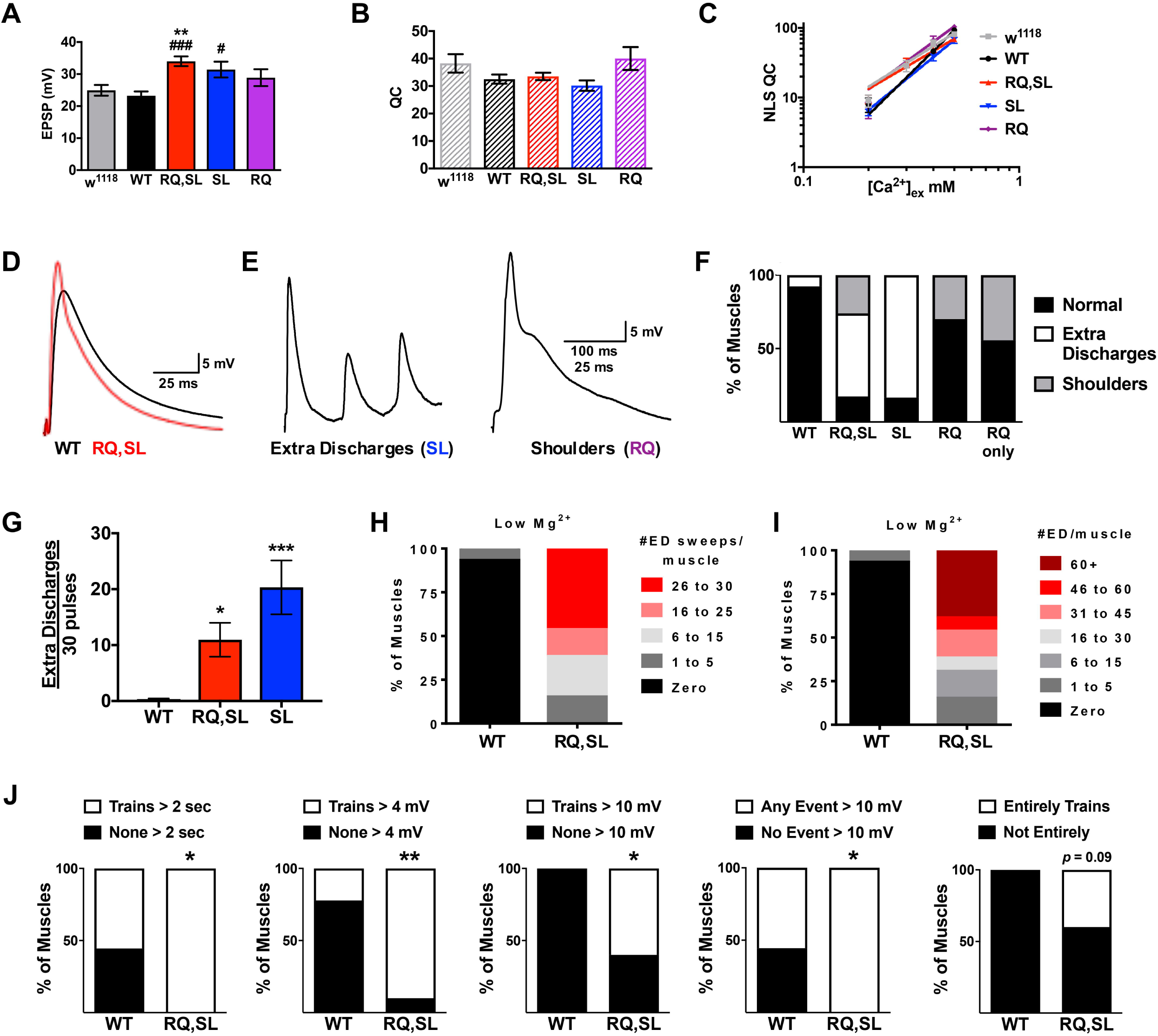
SL- and RQ,SL-expressing NMJs display hyperexcitability in evoked neurotransmission. **(A)** Average EPSP amplitudes at 0.4 mM [Ca^2+^]_e_ for non-transgenic control (*w^1118^*) or Cac-GFP-expressing lines (** *p* < 0.01 by one-way ANOVA with Tukey’s post-hoc vs. *w^1118^*; or # *p* < 0.05 and ### *p* < 0.001 vs. WT; *n* ≥ 12 for all genotypes). **(B)** Average quantal content (QC, estimated as EPSP/mEPSP) at 0.4 mM [Ca^2+^]_e_ (*p* > 0.15 by one-way ANOVA with Tukey’s post-hoc for all genotypes, compared to both *w^1118^* and WT controls). **(C)** Log-log plots of extracellular calcium concentration vs. QC corrected for non-linear summation (NLS QC). There are no statistically significant differences in calcium cooperativity between genotypes (*p* = 0.16, linear regression analysis). **(D, E)** Example electrophysiological traces of **(D)** normal and **(E)** abnormal EPSP waveforms. **(F)** Effect of genotype on EPSP waveforms in response to 30 presynaptic pulses. “RQ only” signifies larvae with a null endogenous *cac* mutation rescued to viability by the RQ-expressing transgene. **(G)** Effect of genotype on number of extra discharges observed per 30 presynaptic pulses (* *p* < 0.05 and *** *p* < 0.001 vs. WT by one-way Kruskal-Wallis ANOVA with Dunn’s post-hoc). **(H)** Penetrance and **(I)** severity of RQ,SL-associated extra discharge waveform dysfunction in low extracellular Mg^2+^ (6 mM). **(J)** NMJ recordings of 2 min spontaneous neurotransmission with an intact CNS. Measurements assessed: continuous trains of spontaneous activity > 2 sec in duration at any point in the recording; trains with postsynaptic events > 4 mV; trains with postsynaptic events > 10 mV; any observed postsynaptic event (trains or not) > 10 mV; any recording that was continuous trains of throughout (*n* = 9 for WT, *n* = 10 for RQ,SL; * *p* < 0.05, ** *p* < 0.01 by Fisher’s Exact Test). All genotypes abbreviated (WT, SL, RQ, RQ,SL) are *elaV(C155)-Gal4/Y; UAS-cac-eGFP^(X)^/+* or *w^1118^* for non-transgenic wild type. Data bars represent the average value and error bars the SEM.

We noted that the EPSP waveforms of RQ, SL, and RQ,SL animals were sometimes abnormal (Figs. 4D, E). In addition to increases in EPSP amplitude (Fig. 4D), we observed two distinct EPSP waveform phenotypes: 1) ‘extra discharges’ (“ED”), in which supernumerary spiking events occurred during the decay phase of the EPSP waveform (Fig. 4E, left); and 2) ‘shoulders,’ in which there was an extended discharge during the decay phase of the EPSP (Fig. 4E, right), causing a discontinuity in the decay. These phenotypes were somewhat reminiscent of a broadening of the end-plate potential previously reported at the NMJs of SL knock-in mice (11). The SL-expressing NMJs produced only the extra discharge type of abnormal waveform, whereas the RQ-expressing NMJs produced only the shoulder form (Figs. 4E, F). Consistent with both mutations being present in the RQ,SL line, those NMJs exhibited both types of abnormal waveform (Fig. 4F).

We were also able to generate “RQ only” animals – functional null X-ray *cac^HC129^* mutant (35) larvae rescued to viability by *elaV(C155)-Gal4-*driven neuronal expression of the RQ transgene. The *cac^HC129^* allele works well for this type of genetic maneuver (22, 28), eliminating endogenous *cac* gene expression, while adding back transgenic *cac*. In the case of “RQ only”, the waveform dysfunction closely matched that shown by the RQ-expressing NMJs (Fig. 4F) – i.e. a shoulder waveform phenotype was present. We were unable to generate “SL only” or “RQ,SL only” animals, possibly due to deleterious gains of function from the SL mutation.

We assessed the severity of the extra discharge phenotype by counting the number of extra discharge events per 30 evoked pulses (30 recording sweeps at 1 Hz per NMJ). Quantification confirmed that SL- and RQ,SL-expressing NMJs were highly dysfunctional, suggesting neuronal hyperexcitability (Fig. 4G). A previous study in Drosophila demonstrated that higher levels of magnesium in the recording saline can mask hyperexcitability of neurons (36). Therefore, we conducted additional WT and RQ,SL recordings in saline with lowered [MgCl_2_] (6 mM vs. 10 mM for normal saline, see Materials and Methods). RQ,SL-expressing NMJs displayed extreme dysfunction in low MgCl_2_, both in terms of the percentage of NMJs that produced any supernumerary discharges (100%, Fig. 4H) and the number of extra discharges counted per 30 presynaptic pulses (Fig. 4I). By contrast, WT-expressing NMJs showed almost no such dysfunction (Figs. 4H, I).

Finally, we conducted an additional series of recordings in normal saline, this time with the larval CNS left intact to check if the hyperexcitability might reflect an *in vivo* state for Drosophila larvae. With this experimental maneuver, it was possible to discern “native circuit” differences between WT- and RQ,SL-expressing animals. The “CNS intact” condition resulted in trains of spontaneous activity. Compared to WT, the RQ,SL-expressing NMJs displayed a high degree of spontaneous activity, marked by rapid, continuous large pulses (Fig. 4J; see several measures and explanation in legend).

In conclusion, SL- and RQ,SL-expressing NMJs displayed evoked gain-of-function phenotypes consistent with prior mammalian FHM1 mutant analyses. By contrast, RQ-expressing NMJs only displayed a mild gain-of-function shoulder phenotype.

### SL- and RQ,SL-expressing NMJs show enhanced spontaneous miniature EPSPs with respect to both amplitude and frequency

Mammalian models of FHM1 show dysfunctional spontaneous neurotransmission (10, 11). We extended our electrophysiological analyses at the Drosophila NMJ to quantal neurotransmission. We observed a striking phenotype: for SL- and RQ,SL-expressing NMJs, there was an enhancement in both amplitude and frequency of spontaneous miniature EPSPs (mEPSPs) (Figs. 5A-E, Table 2). By contrast, neither an increase in spontaneous mEPSP amplitude nor mEPSP frequency were observed for RQ- or WT-expressing NMJs compared to non-transgenic *w^1118^* controls (Figs. 5B-E, Table 2).

**Figure 5:**
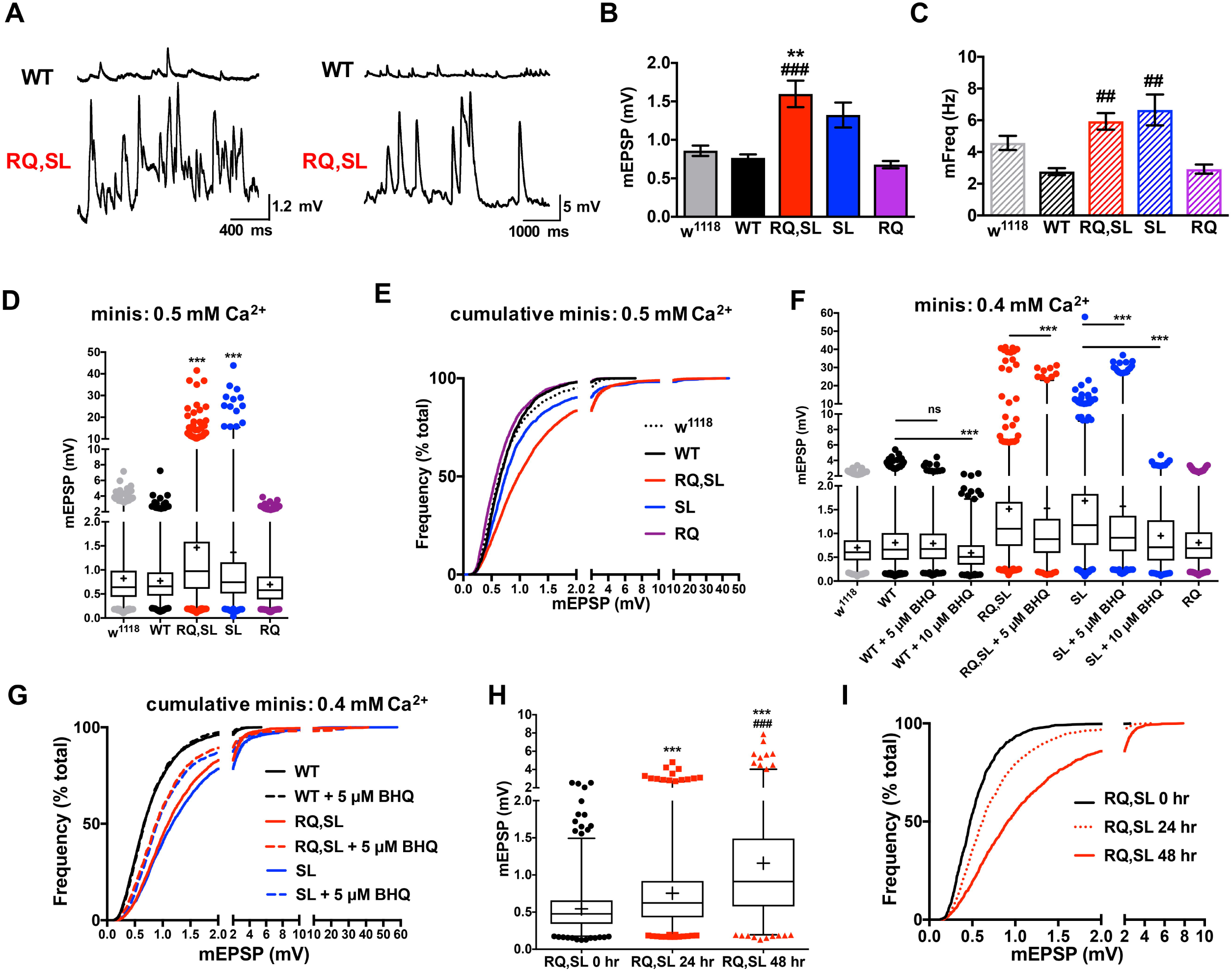
SL- and RQ,SL-expressing NMJs have enhanced mEPSPs. **(A)** Electrophysiological traces of spontaneous activity at WT- and RQ,SL-expressing NMJs. Example traces with two different scales show variable severity of spontaneous neurotransmission phenotypes, in terms of frequency severity (left) or amplitude severity (right). **(B)** Effects of genotype on average spontaneous mEPSP amplitude (** *p* < 0.01 vs. *w^1118^* by one-way ANOVA with Tukey’s post-hoc; ### *p* < 0.001 vs. WT by one-way ANOVA with Tukey’s post-hoc. **(C)** Effects of genotype on spontaneous mEPSP frequency. ## *p* < 0.01 vs. WT by one-way ANOVA with Tukey’s post-hoc; *n* ≥ 12 NMJs, all genotypes. **(D)** Box and whisker plots of mEPSP amplitude range at 0.5 mM extracellular [Ca^2+^]. Box denotes 25^th^-75^th^ percentile; line denotes median; + sign denotes average; whiskers range from 1^st^-99^th^ percentile; individual data points outside the 1^st^ and 99^th^ percentiles are plotted; (*** *p* < 0.001 by Kruskal-Wallis ANOVA with Dunn’s post-hoc vs. either *w^1118^* or WT; *n* > 1400 mEPSPs for each genotype). **(E)** Cumulative probability histogram of the data in (D) showing a marked rightward shift in mEPSP amplitudes for SL-and RQ,SL-expressing NMJs. **(F)** Box and whisker plot (as in (D)) of mEPSP amplitude at 0.4 mM extracellular [Ca^2+^] with and without the Ca_V_2.1 channel modifier BHQ (*** *p* < 0.001 by Kruskal-Wallis ANOVA with Dunn’s post-hoc vs. identical genotype +/- BHQ; *n* > 985 mEPSPs for each genotype). **(G)** Cumulative probability histogram of a subset of data in (F). 5 μM BHQ causes a partial leftward shift in the distribution of events for SL-and RQ,SL-expressing NMJs while not affecting WT-expressing NMJs. **(H)** Box and whisker plot (as in (D)) of mEPSP amplitude range when expressing the RQ,SL transgene for acute periods of developmental time (*** *p* < 0.001 by Kruskal-Wallis ANOVA with Dunn’s post-hoc vs. RQ,SL 0 hr., ### *p* < 0.001 by vs. RQ,SL 24 hr.; *n* > 1095 mEPSPs for each genotype). **(I)** Cumulative probability histogram of the data in (H) showing a rightward shift in mEPSP amplitudes for longer periods of RQ,SL expression.

Since the mutations examined are in a voltage-gated calcium channel, it was important to document electrophysiological behavior at various calcium concentrations. At both 0.5 mM and 0.4 mM extracellular [Ca^2+^], analyses of thousands of individual spontaneous events revealed that increases in spontaneous amplitudes were due to an overall increase in the size distribution of the events at SL- and RQ,SL-expressing NMJs (Figs. 5D-G). Additionally, at both 0.5 mM and 0.4 mM extracellular [Ca^2+^], we noted that the spontaneous events at SL- and RQ,SL-expressing NMJs included a minority of gigantic spontaneous events (10-40 mV) that were never seen in *w^1118^* or WT-expressing controls or in RQ-expressing NMJs (Figs. 5A, D-G). Notably, these gigantic events were seen in the complete absence of presynaptic nerve stimulation in nerves that had already been severed from the central nervous system.

**Table 2.**
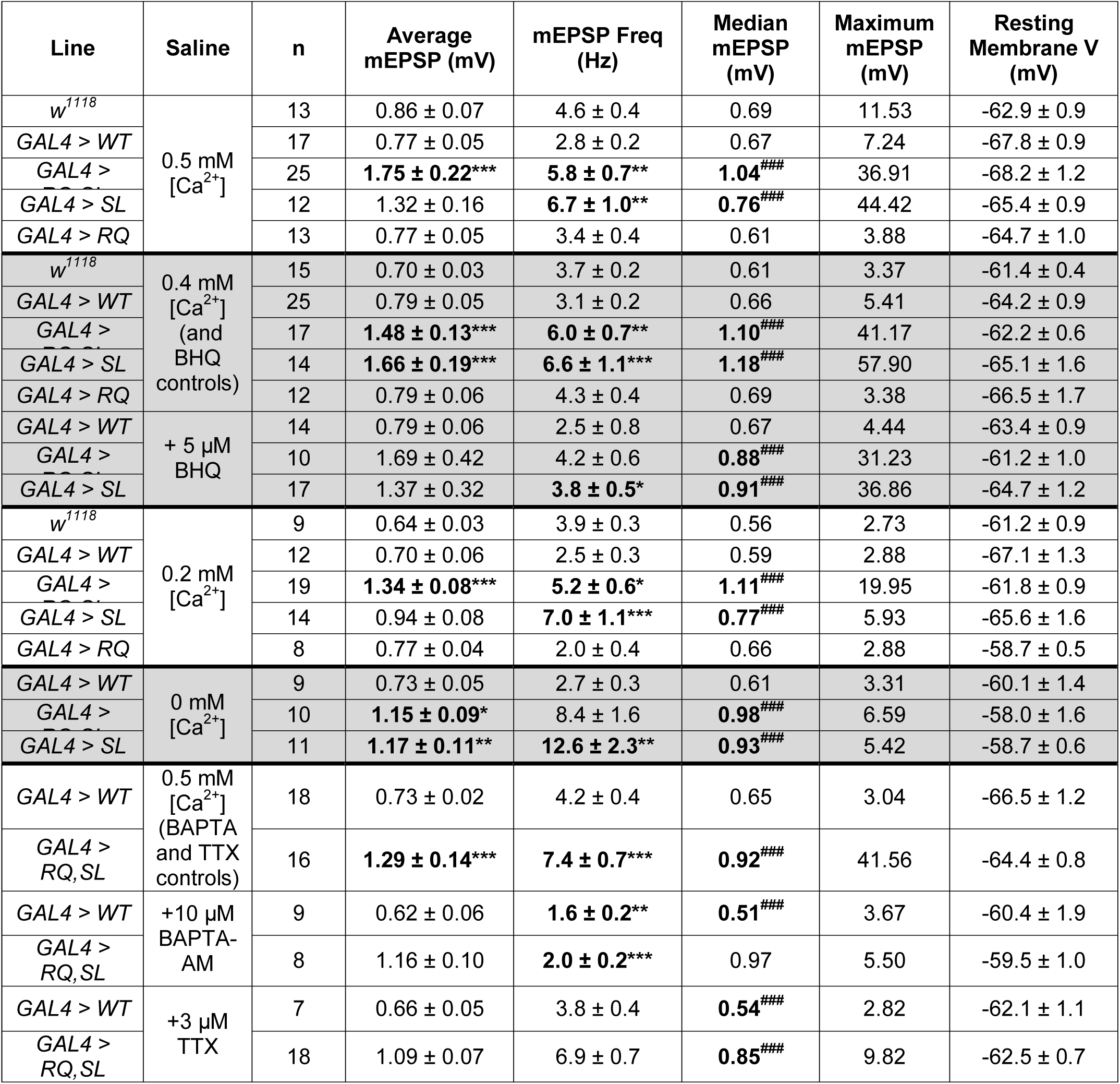
Raw electrophysiological data of selected spontaneous (mEPSP) events. Average mEPSP amplitudes ± SEM and mEPSP frequencies ± SEM for selected conditions. Also given: median mEPSP amplitudes and maximum mEPSP amplitudes achieved for spontaneous events analyzed (~100 per NMJ). *w^1118^* is a non-transgenic wild-type control. WT, RQ, SL, and RQ,SL are shorthand for the indicated *UAS-cac-eGFP* transgene being driven in male progeny presynaptically by the *elaV(C155)-Gal4* driver. These data illustrate differential effects when lowering extracellular [Ca^2+^], chelating Ca^2+^ with BAPTA-AM, or inactivating Na_V_ channels with TTX. Electrophysiological data were analyzed in two ways as average per NMJ and as cumulative distributions. * *p* < 0.05, ** *p* < 0.01, *** *p* < 0.001 vs. control by one-way ANOVA with Tukey’s post-hoc (control is *GAL4 > WT* for most, except in the cases of BHQ, BAPTA-AM, and TTX, in which case the control is the same genotype without treatment). ^###^ *p* < 0.001 vs. control, examining cumulative distributions by Kruskal-Wallis test with Dunn’s post-hoc for multiple comparisons.

It was uncertain if enhanced spontaneous excitability was due to real-time expression of gains of function in Ca_V_2 channel gating kinetics, long-term developmental alterations at the synapse – or if both factors could contribute. We considered altered Ca_V_2 kinetics. It was previously demonstrated that the SL mutation causes complex biophysical alterations to Ca_V_2.1 gating function, both by enhancing voltage-dependent activation (8, 9, 37) and by inhibiting calcium-dependent facilitation (38). Follow-up work showed that the drug 2,5′-di(tertbutyl)-1,4,-benzohydroquinone (BHQ) opposes those effects, reversing SL-induced gains of function (20). As part of the same study, we showed that BHQ restores a form of short-term synaptic plasticity at SL-expressing Drosophila NMJs (20). We extended those prior analyses of BHQ effects on Ca_V_2 gating, this time by examining the distribution of spontaneous events. We found that acute application of 5 μM BHQ was partially effective at reversing the increased size distribution of events for SL- and RQ,SL-expressing NMJs, without changing the distribution of WT events (Figs. 5F, G; Table 2). Notably, 5 μM BHQ did not abolish gigantic events (Figs. 5F, G). Interestingly, a higher concentration of 10 μM BHQ did abolish gigantic events for SL- and RQ,SL-expressing NMJs, but it also significantly decreased the size distribution of WT mEPSPs, which could indicate off-target postsynaptic effects (Fig. 5F). Our BHQ application data are consistent with the idea that that spontaneous neurotransmission gain-of-function phenotypes are driven in part through gating changes at Ca_V_2 channels.

To test if long-term developmental alterations at the synapse could also play a role, we engineered stage-specific *UAS-cac* transgene expression. We utilized the temperature-sensitive *Gal80^TS^/TARGET* system to temporally control expression of the RQ,SL transgene (39). To conduct this experiment, we generated *elaV(C155)-Gal4* >> *UAS-cac-eGFP^RQ,SL^* animals with a ubiquitous *Gal80^TS^* transgene (39). Gal80^TS^ protein halts GAL4-induced gene expression at permissive temperatures (25ºC) but not at restrictive temperatures (29ºC). For our experiment, animals raised at 25ºC throughout life had no discernible spontaneous neurotransmission hyperexcitability (Fig. 5H). By contrast, animals started at 25ºC and shifted to 29ºC for the final 24 or 48 hours before third instar NMJ recording showed progressively more spontaneous hyperexcitability (Figs. 5H, I). This experiment indicates that developmentally regulated expression of gain-of-function Ca_V_2 channel subunits also underlies some of the spontaneous neurotransmission gain-of-function phenotypes.

### Gigantic spontaneous events require extracellular calcium and sodium channel activity

Prior work proposed that mammalian neuronal dysfunction downstream of FHM1 mutations may be calcium-dependent (12). We tested whether the observed effects on quantal size in our model could be calcium-dependent. First, we reduced the extracellular [Ca^2+^] in the recording saline to 0.2 mM. Consistent with classic characterizations of Drosophila NMJ properties (40), low calcium did little to change the distribution of mEPSP size, the median mEPSP size, or the 25^th^-75^th^ percentiles of mEPSP size – all of which remained normal for WT and elevated for SL- and RQ,SL-expressing NMJs (Figs. 6A, B, Table 2). However, lowering extracellular [Ca^2+^] almost completely abrogated gigantic (10-40 mV) spontaneous events at SL- and RQ,SL-expressing NMJs – and it completely eliminated the very largest ones (Figs. 6A, B). This suggested that these gigantic events somehow relied on a sufficient driving force of presynaptic calcium influx – and potentially on spontaneous presynaptic nerve firing.

**Figure 6:**
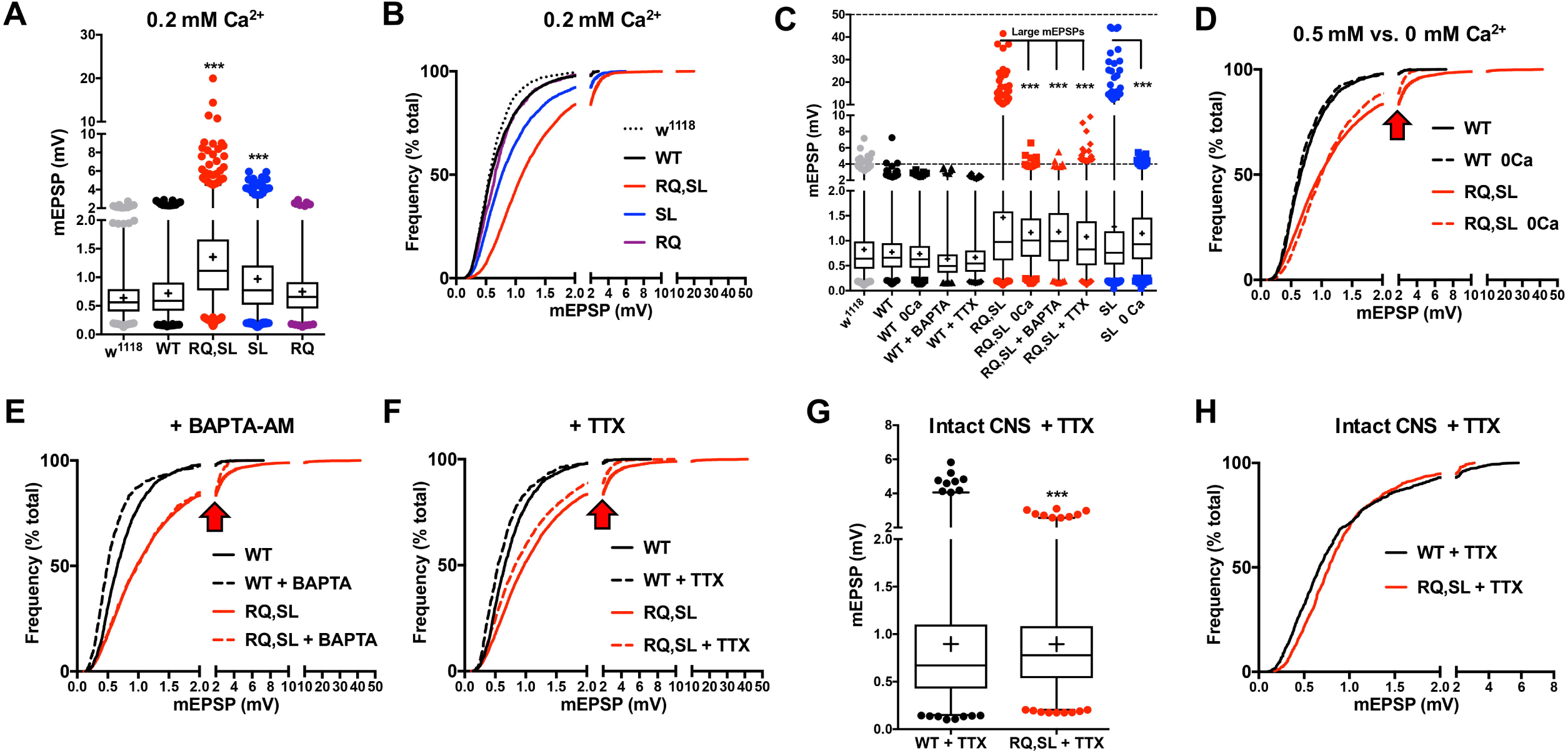
Gigantic spontaneous events vanish in response to diminished Ca^2+^, buffered Ca^2+^, or blocked Na_V_. **(A)** Box and whisker plot of mEPSP amplitudes at 0.2 mM extracellular Ca^2+^. Plot as in Fig. 5 (*** *p* < 0.001 by Kruskal-Wallis ANOVA with Dunn’s post-hoc vs. either *w^1118^* or WT; *n* > 780 mEPSPs for each genotype). **(B)** Cumulative probability histogram of the data in (A) showing a rightward shift in mEPSP amplitudes for SL- and RQ,SL-expressing NMJs – but less so than for 0.5 mM Ca^2+^, with smaller and fewer gigantic events (compare to Figure 5). **(C)** Box and whisker plots demonstrating elimination of gigantic spontaneous events by various manipulations. (*** *p* < 0.001 by Fisher’s exact test examining the incidence of gigantic mEPSPs > 10 mV vs. RQ,SL or SL alone, as appropriate). **(D-F)** Cumulative probability histograms of mEPSP size separately showing the effects of zero extracellular Ca^2+^ (D); application of BAPTA-AM in 0.5 mM Ca^2+^ (E); application of TTX in 0.5 mM Ca^2+^ (F). In each case, the rightward shift in mEPSP size distribution persists due to RQ,SL expression. However, the gigantic spontaneous events are eliminated (see frequency shift at arrowheads). **(G)** Box and whisker plot of spontaneous event amplitudes at 0.5 mM extracellular Ca^2+^ + TTX, with an intact central nervous system. (*** *p* < 0.001 by Mann-Whitney U Test of WT vs. RQ,SL; *n* = 900 mEPSPs for each genotype). **(H)** Cumulative probability histogram of the data in (G).

We extended these analyses by altering the recording saline in three additional ways: 1) zero extracellular calcium; 2) adding the membrane-permeable calcium chelator, 1,2-Bis (2-aminophenoxy) ethane-N,N,N′,N′-tetra acetic acid tetrakis (acetoxymethyl ester) (BAPTA-AM, 10 µM); or 3) adding tetrodotoxin (TTX, 3 μM) to block voltage-gated sodium channels. We compared WT-expressing and RQ,SL-expressing NMJs (and SL-expressing NMJs in the case of zero calcium). All three manipulations produced a similar effect on mEPSP size for the gain-of-function mutants: an elimination of gigantic spontaneous events, but a persistence of overall elevated mEPSP size (Figs. 6C-F, Table 2). By contrast, these manipulations had little to no effect on the distribution of mEPSP amplitudes at WT-expressing NMJs (Figs. 6C-D, Table 2).

Finally, we recorded spontaneous events in more *in vivo*-like condition, using an intact CNS, without severing the motor nerve. In order to do this, we revisited the intact CNS condition (Fig. 4J) – this time adding TTX to the recording saline (0.5 mM [Ca^2+^]). This left the full network anatomy intact, while quieting spontaneous trains of activity. Under these conditions, the spontaneous event amplitude profile of RQ,SL-expressing NMJs was still larger than that of WT-expressing NMJs – and as expected, there were no gigantic events (Figs. 6G, H). Interestingly, however, the difference between WT-expressing NMJs and RQ,SL-expressing NMJs was muted (Figs. 6G, H; compare to Figs. 5D, E). These data suggest that in living animals, network effects could potentially influence the spontaneous gain-of-function activity.

### Large spontaneous events are due to multi-vesicular release

The presence of gigantic spontaneous mEPSPs that were sensitive to low calcium, calcium chelation, and TTX treatment suggested the possibility of spontaneous multi-vesicular release at SL- and RQ,SL-expressing NMJs. If this were true, traditional analysis of spontaneous mEPSPs would result in an overestimation of average quantal size (Fig. 5B) and underestimation of average QC (Fig. 4B) for SL-and RQ,SL-expressing NMJs.

We utilized the method of failures to better resolve questions about quantal size and QC. At very low concentrations of extracellular calcium, synapses like the NMJ are essentially limited to a one-or-none evoked response in which stimulation of the presynaptic nerve either leads to the release of a single vesicle or fails to release any vesicles (41). By conducting failure analyses, it is possible to measure the distribution of quantal events and also to estimate QC in a way that eliminates confounds of higher concentrations of calcium. First, we conducted failure analysis recordings at 0.14 mM [Ca^2+^]_e_ for WT-, RQ,SL-, and SL-expressing NMJs (Figs. 7A-C). For this condition, the evoked events for SL- and RQ,SL-expressing NMJs were far larger on average than those observed WT-expressing NMJs (Fig. 7C – EPSP). This was due to a large proportion of events of > 2 mV for the SL- and RQ,SL-expressing conditions (compare Figs. 7A, B). Furthermore, even in this low level of extracellular Ca^2+^, many of the RQ,SL and SL events represented multi-vesicular release rather than the release of a single large vesicle. We calculated values of QC of > 2 for both mutant conditions at 0.14 mM [Ca^2+^]_e_ (QC = *m* = ln[(# trials)/(# failures)] (42)) (Fig. 7C).

**Figure 7:**
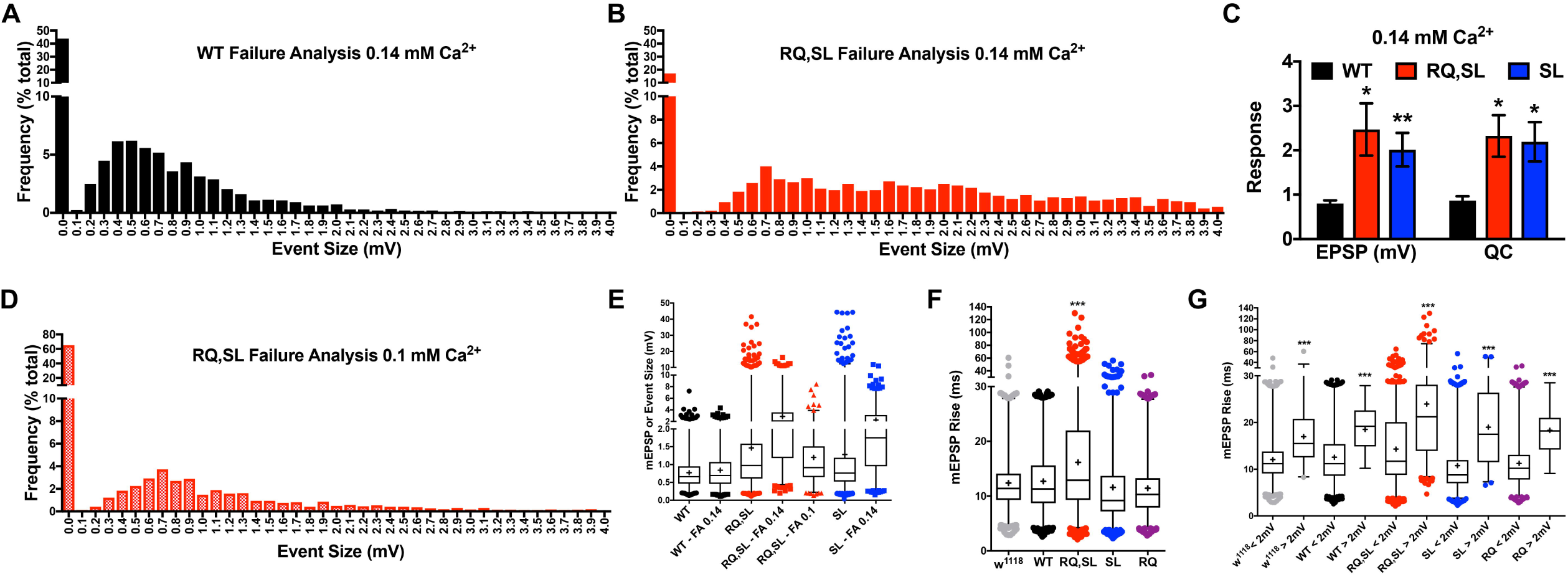
Failure analysis: SL- and RQ,SL-expressing NMJs show elevated release probability at very low extracellular calcium. **(A-B)** Frequencies of evoked amplitudes at very low extracellular Ca^2+^ (0.14 mM) for **(A)** WT-expressing NMJs and **(B)** RQ,SL-expressing NMJs. For the RQ,SL-expressing NMJs, there is a clear rightward shift in the size distribution of RQ,SL-expressing events, as well as a marked decrease in the frequency of failures (categorized as 0 mV events). **(C)** For WT-, SL-, and RQ,SL-expressing NMJs, the average EPSP size for successfully evoked events, as well as estimated QC by failure analyses (0.14 mM Ca^2+^) (* *p* < 0.05; ** *p* < 0.01 by one-way ANOVA with Tukey’s post-hoc compared to WT). **(D)** Further lowering extracellular Ca^2+^ (0.1 mM) for RQ,SL reveals a leftward shift in size distribution and an increase in failure percentage compared to (B). **(E)** Box and whisker data are presented as in Figs. 5 and 6 – this time showing the size distributions of spontaneous mEPSP events (WT, SL, RQ,SL), as well as failure analysis (FA) evoked events for the same genotypes (failures excluded). **(F, G)** Box and whisker plots for mEPSP rise times (0.5 mM Ca^2+^, see Fig. 5D) show a significant increase only for RQ,SL-expressing NMJs (F), as well as a dramatic slowdown for events > 2 mV in size, regardless of genotype (G).

To test if lower calcium could generate a leftward shift in event size, we applied a more restrictive condition of 0.1 mM [Ca^2+^]_e_ to RQ,SL-expressing NMJs. At 0.1 mM [Ca^2+^]_e_ the proportion of failures was very high for RQ,SL-expressing NMJs, with events over 4 mV all but absent, and events greater than 1.5 mV also less prevalent (Fig. 7D). The first peak in the distribution of events, which is reflective of single vesicle size (42), was centered near 0.7 mV (Fig. 7D), a value consistent with single-vesicle responses of normal size for the Drosophila NMJ (40). Together, these data suggested that the observed large events at SL- and RQ,SL-expressing NMJs – regardless of whether spontaneous or failure analysis-evoked – were likely due to multi-vesicular release (see Fig. 7E, spontaneous and failure analyses distributions side-by-side).

If larger spontaneous events are multi-vesicular (or at the very least include a proportion of multi-vesicular events), this property should also be reflected in slowed spontaneous event rise time kinetics. We analyzed the rise time kinetics of several thousand spontaneous events for *w^1118^*, WT-, RQ-, SL-, and RQ,SL-expressing NMJs. Average rise times were slowed only for RQ,SL-expressing NMJs (Fig. 7F). However, the rise times for larger events were markedly slower for *all* genotypes, not just RQ,SL (Fig. 7G). For SL- and RQ,SL-expressing NMJs there was a much larger proportion of such events. Collectively, our data suggest that large events (> 2 mV) include several that are multi-vesicular.

### PLCβ loss genetically suppresses spontaneous excitability

For SL- and RQ,SL-expressing NMJs, we hypothesized that specific cellular cues could dictate the various electrophysiological phenotypes we documented: multi-vesicular quantal events, gigantic TTX-sensitive spontaneous events, and enhanced NMJ excitability. We inquired as to what the molecular nature of those cues might be. Our experiments indicated that intracellular calcium or intracellular calcium signaling processes might be important (Fig. 6). Additionally, recent data from the mouse calyx of Held demonstrated that S218L knock-in synapses have enhanced resting intracellular calcium (12). We hypothesized that altered intracellular calcium signaling or handling could impact myriad intracellular signals and investigated which signaling pathways might be relevant. This line of inquiry spurred a genetic approach examining regulators of intracellular calcium to test if inhibition of any of these factors may influence gain-of-function Ca_V_2 phenotypes at the synapse (Fig. 8A). We sought to identify suppressors capable of reversing gains of Ca_V_2 function caused by the SL and RQ,SL transgenes.

**Figure 8:**
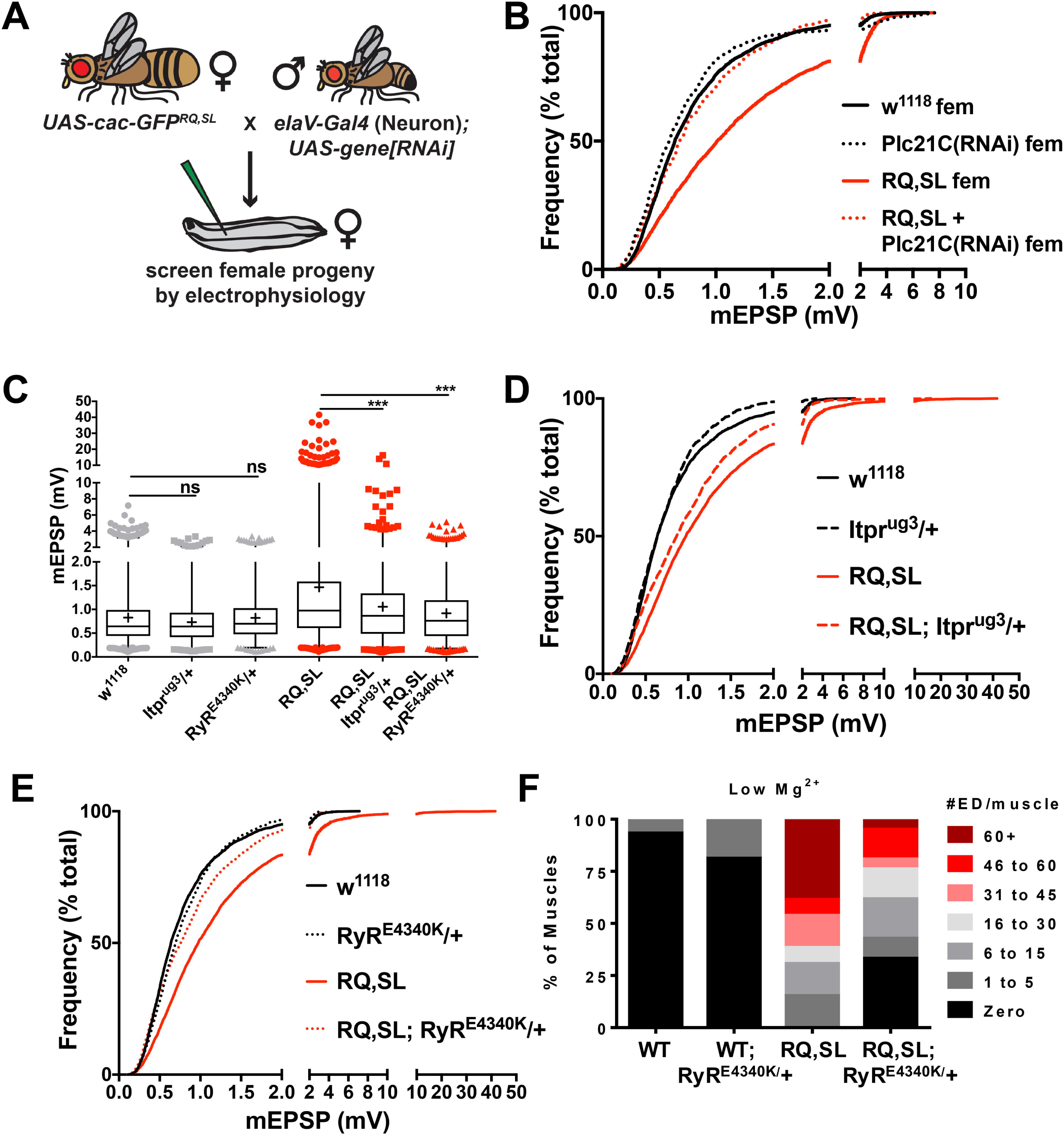
Inhibition of an intracellular Ca^2+^ release pathway dampens gain-of-function phenotypes associated with FHM1-mimicking mutations. **(A)** Schematic of an RNA interference (RNAi)-based approach to identify suppressors of gain-of-function electrophysiological phenotypes. **(B)** Knockdown of *Plc21C* gene function reverses the increase in spontaneous mEPSP amplitude elicited by RQ,SL expression. **(C)** Box and whisker plots (as before) and **(D, E)** cumulative probability histograms (as before) demonstrate that heterozygous, loss-of-function point mutations in genes encoding the IP_3_ receptor (*Itpr^ug3^/+*) and the Ryanodine receptor (*RyR^E4340K^/+*) also significantly diminish the gain-of-function spontaneous mEPSP phenotypes in RQ,SL-expressing NMJs. *** *p* < 0.001 by Kruskal-Wallis ANOVA with Dunn’s multiple comparisons test vs. RQ,SL alone. **(F)** The *RyR^E4340K^/+* condition also diminishes evoked EPSP hyperexcitability phenotypes in a RQ,SL-expressing background (# of extra discharges [ED] per muscle – see also Figure 4).

Prior studies of Drosophila NMJ homeostatic synaptic plasticity, which involves the potentiation of Ca_V_2 function, suggested some possible candidate molecules (43, 44). Additionally, we previously showed that the Drosophila PLCβ homolog *phospholipase-C at 21C* (*Plc21C*) is necessary for this same neuronal homeostatic potentiation mechanism (45). Plc21C is one of two Drosophila Phospholipase-Cβ (PLCβ) family members, and is expressed in the nervous system (46). Canonically, PLCβ proteins cleave phosphatidylinositol 4,5-bisphosphate (PIP_2_) to generate soluble inositol triphosphate (IP_3_), as well as membrane-bound diacylglycerol (DAG). These signaling factors influence synaptic transmission in a variety of ways, including direct modulation of Ca_V_2 (47), and they have been shown to act at several synapses, including the NMJ (48–53).

We targeted *Plc21C* gene expression in neurons with a previously verified *UAS-Plc21C(RNAi)* construct, *Plc21C^GD11359^* (45, 54). Compared to the NMJs of *w^1118^* and WT controls, those in which only *Plc21C* had been knocked down presynaptically exhibited no discernable baseline changes in mEPSP size (Fig. 8B, Table 3) – or as previously documented, EPSP size, or QC (45). By contrast, in RQ,SL-expressing NMJs such *Plc21C* knockdown alleviated aspects of NMJ hyperexcitability. Specifically, there was a leftward shift in the distribution of spontaneous events (Fig. 8B, Table 3). Interestingly, there was not a significant reversal of the enhanced mEPSP frequency phenotype (Table 3).

**Table 3.** Raw electrophysiological data of spontaneous (mEPSP) events – impairment of intracellular Ca^2+^ release pathway. Average mEPSP amplitudes ± SEM and mEPSP frequencies ± SEM for selected experimental conditions. Also given are the median mEPSP amplitude and the maximum mEPSP amplitudes achieved for all spontaneous events analyzed per genotype (~100 per NMJ). *w^1118^* is a non-transgenic wild-type control. WT and RQ,SL are shorthand for the indicated *UAS-cac-eGFP* transgene being driven in progeny presynaptically by the *elaV(C155)-Gal4* driver. This table illustrates differential effects when impairing an intracellular calcium release signaling pathway through mutation of the *Plc21C*, *itpr*, and *RyR* genes, or through pharmacological application of Xestospongin C or LiCl. Electrophysiological data were analyzed in two ways as average per NMJ and as cumulative distributions. * *p* < 0.05, ** *p* < 0.01, *** *p* < 0.001 vs. control by one-way ANOVA with Tukey’s post-hoc (for all cases, the appropriate control is the same genotype without treatment; some control data are on Table 2). ^#^ *p* < 0.05, ^##^ *p* < 0.01, ^###^ *p* < 0.001 vs. control to examine cumulative distributions by Kruskal-Wallis test with Dunn’s post-hoc.

| Line | Experiment (all 0.5 mM $\text{Ca}^{2+}$) | n | Average mEPSP (mV) | mEPSP Freq (Hz) | Median mEPSP (mV) | Maximum mEPSP (mV) | Resting Membrane V (mV) |
| --- | --- | --- | --- | --- | --- | --- | --- |
| *w<sup>1118</sup>* females | *Plc21C* RNAi baseline | 27 | $0.81 \pm 0.05$ | $4.3 \pm 0.3$ | 0.64 | 7.16 | $-62.2 \pm 0.6$ |
| *Plc21C(RNAi)* males | | 12 | $0.80 \pm 0.06$ | **$2.0 \pm 0.2^{**}$** | 0.61 | 4.50 | $-66.5 \pm 0.9$ |
| *Plc21C(RNAi)* females | | 6 | $0.83 \pm 0.02$ | $3.8 \pm 0.9$ | 0.59 | 7.68 | $-61.6 \pm 0.3$ |
| *GAL4 > RQ,SL* females | suppression of RQ,SL | 19 | $1.28 \pm 0.08$ | $5.8 \pm 0.5$ | 1.01 | 7.62 | $-67.0 \pm 1.2$ |
| *GAL4 > RQ,SL + Plc21C(RNAi)* | | 13 | **$0.78 \pm 0.04^{***}$** | $5.2 \pm 0.6$ | **$0.68^{***}$** | 3.89 | $-64.9 \pm 1.6$ |
| *GAL4; itpr<sup>μg3</sup>/+* | suppression of RQ,SL | 9 | $0.74 \pm 0.04$ | $3.7 \pm 0.2$ | 0.64 | 3.34 | $-61.3 \pm 0.6$ |
| *GAL4 > RQ,SL; itpr<sup>μg3</sup>/+* | | 14 | **$1.05 \pm 0.06^*$** | $6.1 \pm 0.5$ | **$0.86^{***}$** | 16.21 | $-65.8 \pm 0.7$ |
| *GAL4; RyR<sup>E4340K</sup>/+* | suppression of RQ,SL | 13 | $0.83 \pm 0.03$ | $3.3 \pm 0.3$ | 0.70 | 3.38 | $-63.0 \pm 0.5$ |
| *GAL4 > RQ,SL; RyR<sup>E4340K</sup>/+* | | 17 | **$0.91 \pm 0.04^{***}$** | $5.1 \pm 0.8$ | **$0.76^{***}$** | 5.13 | $-62.5 \pm 0.9$ |
| *GAL4 > WT* | XestC and LiCl controls | 19 | $0.80 \pm 0.02$ | $2.3 \pm 0.3$ | 0.71 | 2.59 | $-69.6 \pm 1.1$ |
| *GAL4 > RQ,SL* | | 30 | $1.63 \pm 0.13$ | $6.6 \pm 0.7$ | 1.08 | 51.03 | $-65.8 \pm 0.7$ |
| *GAL4 > WT* | + 5 $\mu\text{M}$ XestC | 7 | $0.89 \pm 0.03$ | $3.9 \pm 0.9$ | 0.78 | 3.80 | $-69.8 \pm 2.2$ |
| *GAL4 > RQ,SL* | | 14 | **$1.14 \pm 0.11^*$** | $6.4 \pm 1.1$ | **$0.81^{***}$** | 30.76 | $-67.2 \pm 1.3$ |
| *GAL4 > WT* | + 10 mM LiCl | 11 | $0.81 \pm 0.03$ | $3.2 \pm 0.4$ | 0.73 | 4.09 | $-66.6 \pm 1.3$ |
| *GAL4 > RQ,SL* | | 12 | **$1.2 \pm 0.06^*$** | $4.1 \pm 0.4$ | **$0.98^{***}$** | 6.34 | $-68.8 \pm 1.7$ |

### IP_3_R and RyR point mutations strongly suppress hyperexcitability

We hypothesized that Plc21C could exert effects on spontaneous neurotransmission via one of several components of its canonical signaling pathway (e.g. PIP_2_, DAG, or IP_3_). Notably, IP_3_ acts through the IP_3_ receptor (IP_3_R), an intracellular calcium channel located on the endoplasmic reticulum (ER). At other model synapses, release of Ca^2+^ from the intracellular stores can promote the release of neurotransmitter-laden vesicles and contribute to the amplitudes of spontaneous events (55–58). Moreover, IP_3_R has been proposed to play a role in spontaneous vesicle release through calcium-induced calcium release (CICR) (59), and increased ER Ca^2+^ release was recently shown to potentiate synaptic transmission at the Drosophila NMJ (60).

We examined the Drosophila IP_3_R gene (*itpr*). Homozygous *itpr* loss-of-function mutations are lethal, so we tested a heterozygous loss-of-function condition. Since IP_3_R clusters consist of multiple units, we hypothesized that we might be able to partially disrupt them through a loss-of-function point mutation, *itpr^ug3^*, a mutant possessing a missense mutation in the IP_3_R ligand-binding domain (61). *itpr^ug3^/+* phenocopied *Plc21C* knockdown at RQ,SL-expressing NMJs: the mEPSP amplitude was partially reduced toward WT levels, and the number of giant, spontaneous events was diminished (Figs. 8C, D, Table 3). Importantly, on its own *itpr^ug3^/+* did not significantly affect the baseline amplitude or distribution of mEPSPs (Fig. 8C, Table 3). Finally, as with the RNAi experiment, the increased mEPSP frequency phenotype was not suppressed (Table 3).

We performed analogous experiments with a Drosophila ryanodine receptor gene (*RyR*) mutation. Tetrameric RyR channels have been reported to contribute to CICR downstream of IP_3_Rs (59). Additionally, gigantic spontaneous miniature potentials at other model synapses are mediated by RyR and rapid expulsion of calcium from presynaptic stores (62–66). We found that the heterozygous *RyR* point mutant *RyR^E4340K^/+* (67) almost completely suppressed the increased average mEPSP amplitude in the RQ,SL-expressing background (Figs. 8C, E, Table 3). Additionally, the gigantic spontaneous events were abrogated (Figs. 8C, E, Table 3). Control recordings showed that *RyR^E4340K^/+* did not affect the baseline amplitude or distribution of mEPSPs (Figs. 8C, E). As with *Plc21C* and *itpr*, impairment of *RyR* function did not significantly suppress the enhanced mEPSP frequency phenotype of RQ,SL-expressing NMJs (Table 3).

Because the *RyR^E4340K^/+* background provided such a strong suppression of spontaneous mEPSP hyperexcitability at RQ,SL-expressing NMJs, we checked if it could also suppress hyperexcitability in the context of evoked excitation. As shown before, when incubated in low extracellular magnesium, 100% of the RQ,SL-expressing NMJs showed a hyperexcitability dysfunction, with high expressivity of extra discharges (Figs. 3H,I; 8F). In a heterozygous *RyR^E4340K^*/+ genetic background, this hyperexcitability phenotype was partially suppressed, in terms of both the penetrance of NMJs with extra evoked discharges and the expressivity of the extra discharge dysfunction at individual NMJs (Fig. 8F). On its own, the *RyR^E4340K^/+* condition shows almost no baseline hyperexcitability phenotype (Fig. 8F).

### Spontaneous mEPSP hyperexcitability can be suppressed pharmacologically

Our data for genetic manipulations affecting Plc21C, IP_3_R, and RyR show that it is possible to attenuate RQ,SL-induced gain-of-function mEPSP amplitude and excitability phenotypes by genetically impairing factors known to promote intracellular Ca^2+^ release. We wondered if pharmacological manipulations could also be effective. We turned to two agents to test this idea: lithium (10mM LiCl in larval food) and Xestospongin C (5 μM in recording saline). Chronic exposure to lithium inhibits inositol monophosphate phosphatase, eventually resulting in a disruption of the recycling process that generates PIP_2_ (68, 69). Xestospongin C has been previously characterized as a membrane-permeable inhibitor of IP_3_ receptors (70, 71). Either chronically feeding larvae LiCl or applying Xestospongin C to the recording bath caused a significant leftward shift in the overall size distribution of spontaneous amplitudes (Figs. 9A-C), reminiscent of the effects observed for *Plc21C*, *itpr*, and *RyR* losses of function. The acute Xestospongin C application seemed to exert a stronger suppression effect in this regard, while the chronic LiCl application exerted a stronger suppression of the gigantic spontaneous events (Figs. 9A-C, Table 3). Notably, neither pharmacological manipulation diminished baseline spontaneous neurotransmission in WT-expressing control NMJs, nor did either manipulation significantly suppress the elevated mEPSP frequency phenotype for RQ,SL-expressing NMJs (Figs 9A-C, Table 3).

**Figure 9:**
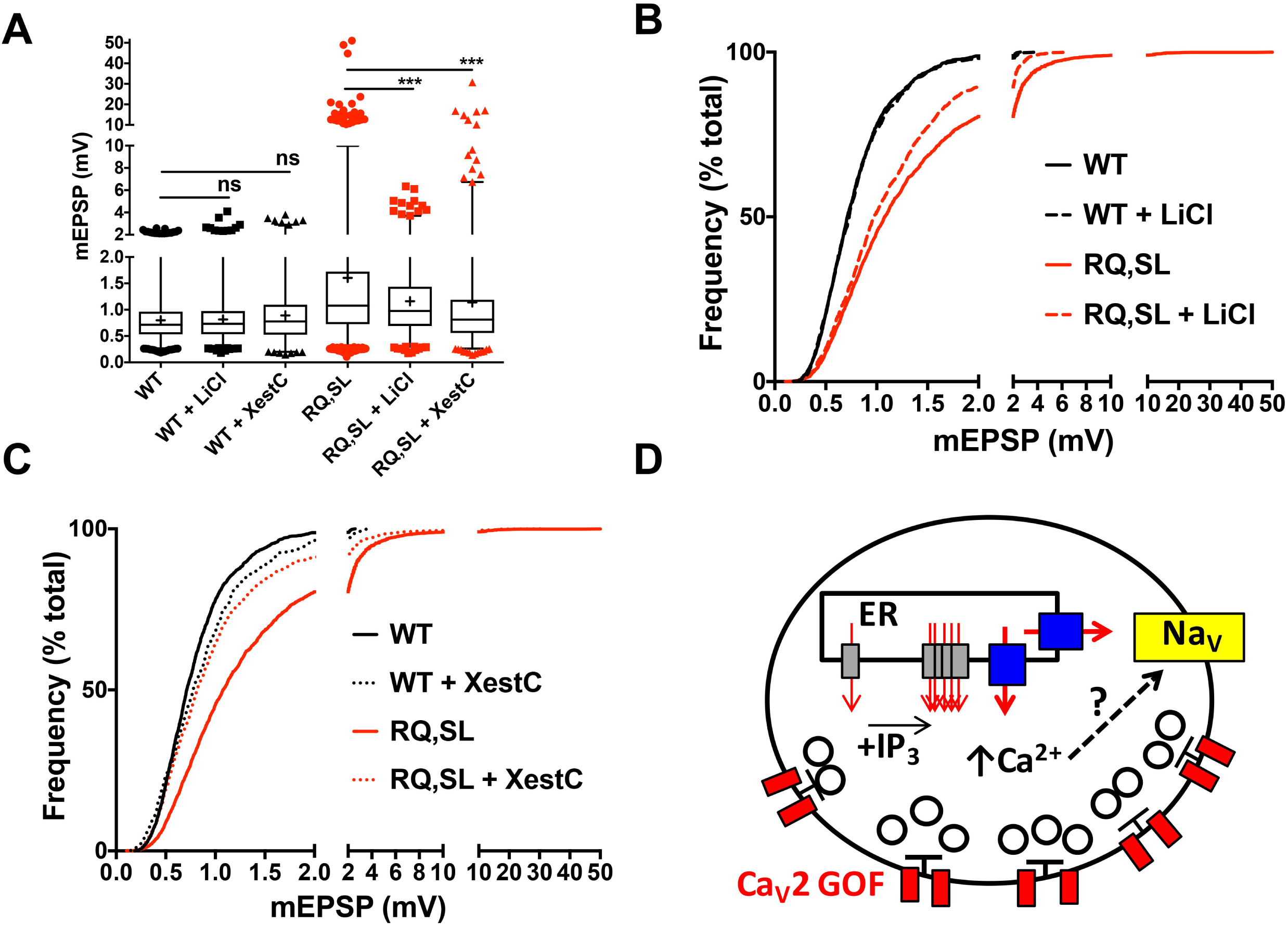
Pharmacological inhibition of intracellular Ca^2+^ release dampens gain-of-function phenotypes associated with FHM1-mimicking mutations. **(A-C)** Data displayed and analyzed as before. Box and whisker plots (A) and cumulative probability histograms (B, C) demonstrate that acute application of either LiCl (to block PIP_2_ recycling) or Xestospongin C (to block IP_3_ receptors) both suppress the gain-of-function spontaneous mEPSP phenotypes in RQ,SL-expressing NMJs. *** *p* < 0.001 by Kruskal-Wallis ANOVA with Dunn’s multiple comparisons test vs. RQ,SL alone. **(D)** Cartoon model depicting neuronal components implicated in this study of regulating neurophysiology downstream of migraine-mimicking amino-acid substitutions. Red – Ca_V_2 channels; gray – IP_3_ receptors; blue – Ryanodine receptors; yellow – Na_V_ channels.

### Mutations targeting intracellular calcium release signaling can exacerbate lethality

Since genetic mutations that target intracellular calcium release ameliorate hyperexcitability phenotypes, we reasoned that the same (or similar) mutations might ameliorate the lethality phenotypes associated with expressing the RQ,SL transgene. We conducted lethality test crosses and progeny counts in a similar manner as before (Table 1). This time, we crossed females bearing the *UAS-cac-eGFP^RQ,SL^* transgene to males carrying both the *elaV(C155)-Gal4* driver and a collection of loss-of-function genetic manipulations on *Drosophila melanogaster* Chromosome II for the *Plc21C*, *RyR*, or *Gq* genes. In addition to *Plc21C* and *RyR*, we chose *Gq* because canonical PLCβ signaling is downstream of Gαq function. Our prior work showed that *Plc21C* and *Gq* play a role in the maintenance of homeostatic plasticity at the NMJ (45). The hypothesis to test was that female progeny carrying the driver, the RQ,SL transgene, and the intracellular calcium release manipulation could have improved viability versus to female progeny carrying only the driver and the RQ,SL transgene. Male progeny siblings would not carry the driver – and would therefore not express the RQ,SL transgene – and could be used to control for parameters affecting lethality, independent of the RQ,SL transgene.

As expected, female progeny carrying the driver, the RQ,SL transgene, and no balancer chromosome had reduced viability compared to their male sibling counterparts (Table 4; see “+”). However, introducing heterozygous loss-of-function manipulations affecting *Plc21C*, *RyR*, and *Gq* did not ameliorate this phenotype. Surprisingly, those manipulations almost always further reduced viability, often strongly (Table 4). The effect was particularly strong for all *Plc21C* and *Gq* loss-of-function conditions examined (Table 4). For *RyR*, the effect was strong only for the *RyR^16^* deletion allele (Table 4). Heterozygous *RyR* point mutant manipulations did not further enhance lethality in a statistically significant way, but they did not ameliorate the lethality phenotype either (Table 4). No manipulation examined resulted in significantly higher male lethality, compared to control males (Table 4). These results highlight the fact that molecular manipulations can have a salubrious effect in one context (synapse excitability) and an exacerbating effect in another (viability). This is true in our Drosophila system but possibly in other systems as well.

**Table 4.**
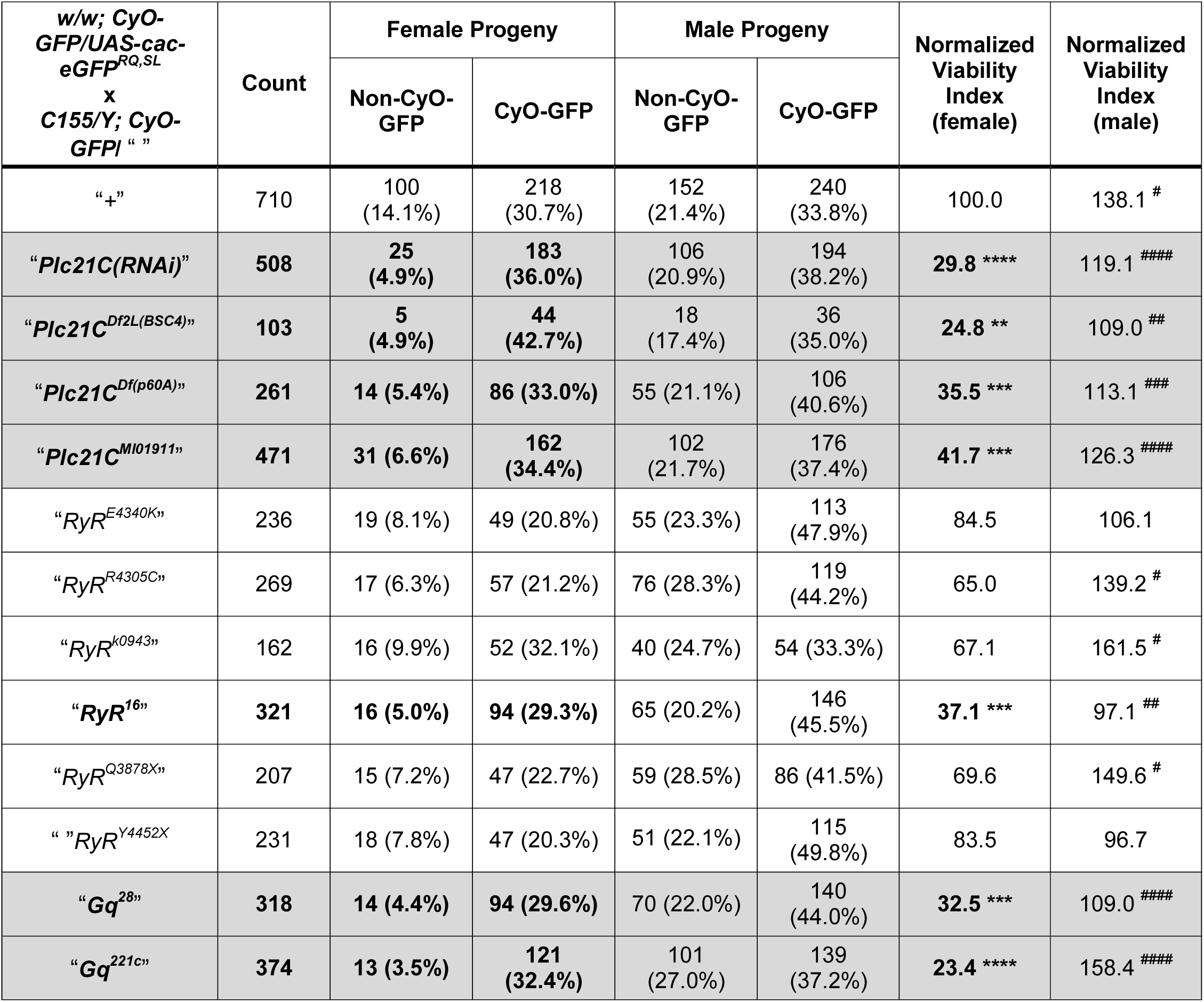
Loss-of-function mutations in an intracellular Ca^2+^ store release pathway enhance adult lethality phenotypes. Viability enhancement/suppression test crosses were performed utilizing *w/w; CyO-GFP/UAS-cac-eGFP^RQ,SL^* virgin females x *elaV(C155-Gal4)/Y; CyO-GFP/* “*mutant or UAS-RNAi or +*” males. Balancer or non-Balancer (CyO-GFP and non-CyO-GFP) female and male progeny were counted. Raw progeny counts and relative proportions are shown. Changes in the proportion of non-CyO-GFP female progeny acquired could indicate a suppression or enhancement effect on viability. A normalized viability index number was set = (proportion of non-CyO-GFP females for “genotype”)/(proportion of non-CyO-GFP females for “+”). In all cases for losses of function of *Plc21C*, *RyR*, and *Gq* gene function, the normalized viability index decreased numerically, but not always to a statistically significant degree. Fisher’s exact tests were performed for each cross to test for differences in female to male CyO-GFP:non-CyO-GFP ratios. This type of analysis controlled for any lethality caused by the genetic manipulation itself. For female progeny ratios, * *p* < 0.05, ** *p* < 0.01, *** *p* < 0.001, **** *p* < 0.0001 vs. *“*+” by Fisher’s exact test between crosses. Within a cross, ^#^ *p* < 0.05, ^##^ *p* < 0.01, ^###^ *p* < 0.001, ^####^ *p* < 0.0001 for male vs. female viable progeny ratios. Most crosses meet a significant threshold by this latter criterion because the *RQ,SL* transgene causes lethality itself, and it is only expressed in females.

## DISCUSSION

We generated fruit flies designed to mimic the effects of FHM1-inducing Ca_V_2.1 channel mutants, R192Q and S218L. Flies expressing the SL and RQ,SL transgenes for Drosophila Ca_V_2/Cacophony displayed overt phenotypes, including reduced viability (Fig. 1). They also displayed synaptic phenotypes, including enhanced evoked excitability (Fig. 4), stark increases in quantal size and frequency (Fig. 5), giant, spontaneous, sodium channel-dependent events (Figs. 5, 6), and enhanced probability of release at very low calcium (Fig. 7). All of these neurotransmission phenotypes occurred without major alterations in active zone localization or overall synaptic architecture (Figs. 2, 3). RQ-expressing NMJs had only a mild phenotype: EPSP discharges with extended, shoulder-like waveforms (Fig. 4). Genetic knockdown of Drosophila PLCβ or genetic mutations affecting the receptors that gate intracellular calcium stores (IP_3_ receptor and Ryanodine receptor) partially alleviated some of the electrophysiological phenotypes (Fig. 8), as did pharmacological manipulations targeting the same processes (Fig. 9). These results suggest that intracellular Ca^2+^ signaling through IP_3_ receptors and Ryanodine receptors could influence physiological dysfunction in a gain-of-function Ca_V_2 background (Fig. 9D). Additionally, given the ability of TTX to block gigantic spontaneous events – and given our ability to quiet that phenotype through genetic and pharmacological means – impairment of the IP_3_ Receptor/Ryanodine Receptor pathway may limit spontaneous neuronal firing by as-yet undetermined mechanisms (Fig. 9D).

### Similarities between fly mutations and FHM1-causing human mutations

*Evoked Neurotransmission*. Our discovery that SL- and RQ,SL-expressing Drosophila NMJs displayed increased evoked excitation, especially at low [Ca^2+^]_e_ (Figs. 4, 7) (20), was consistent with findings from diaphragm NMJs in SL knock-in mice (11). In that context, the end-plate potential (EPP) amplitudes were significantly increased at low levels of calcium (0.2 mM) but did not differ from those at wild-type NMJs at physiological calcium (2 mM) (11). Interestingly, at the SL knock-in calyx of Held, excitatory postsynaptic currents (EPSCs) were increased, but this effect was most pronounced at high levels of [Ca^2+^]_e_ (12). The EPSP discharges caused by expression of the SL-containing transgenic constructs in flies (Fig. 4) were reminiscent of the EPP broadening at SL knock-in NMJs (11). Finally, the severity of the dysfunction in the Drosophila NMJ waveform in the context of decreased extracellular magnesium (6 mM) (Figs. 4, 8) was consistent with a marked increase in calcium current in response to long action potential waveforms in calyces of Held expressing the RQ or SL mutant protein (12, 14).

*Enhanced Quantal Frequency*. The enhanced mEPSP frequency at SL-and RQ,SL-expressing Drosophila NMJs (Fig. 5, Table 1) was reminiscent of observations in prior FHM1 studies. In the RQ and SL knock-in mice, the NMJs exhibited significant increases in the frequency of mEPPs (9–11). In principle, this spontaneous activity could correlate with a buildup of intracellular calcium or a change in intracellular calcium dynamics. In support of this view, at the calyx of Held in SL knock-in mice the frequency of spontaneous mEPSCs was enhanced and resting [Ca^2+^]_i_ was elevated (12). In that case, the increase in quantal frequency was partially reversed by adding the cell-permeable calcium chelator EGTA-AM (12).

Evidence from several model synapses suggests that Ca_V_2 channels can play a prominent role in spontaneous release. In granule cells of the hippocampus, stochastic activity of Ca_V_2.2 (N-type) channels potentiates spontaneous miniature events, and the application of either BAPTA-AM or EGTA-AM is sufficient to inhibit them (72). Other studies have demonstrated that P/Q-, N-, and R-type calcium channels also promote spontaneous release (73). Notably, the differences in the spontaneous miniature phenotype between mice harboring the SL and RQ knock-in substitutions, or fruit flies expressing mimicking substitutions, suggest that the differences in cellular outcomes occur downstream of the Ca_v_2 channel. This highlights a need for genetic approaches to uncover pathways that might contribute to the divergent phenotypes, as well those that are shared.

### Differences between fly mutations and FHM1-causing human mutations

*Quantal amplitudes*. FHM1 mutations have been shown to enhance spontaneous miniature quantal release frequency in other systems (8–11), but there has been no report of increases in spontaneous miniature quantal size due to these mutations. In theory, an increase in the amplitude of mEPSP events at the Drosophila NMJ could be explained by an alteration to the expression and localization of postsynaptic proteins. Yet immunostaining of postsynaptic markers showed only a slight increase in postsynaptic glutamate receptor clustering (Fig. 3G). Instead, a combination of quantal analyses (Figs. 5–7) points to alterations to the nature of spontaneous, presynaptic vesicle release – namely, that a certain percentage of quantal events in SL- and RQ,SL-expressing NMJs are multi-vesicular.

Why do SL- and RQ,SL-expressing NMJs in Drosophila show spontaneous multi-vesicular release? The synaptic preparation examined is likely critical. Evidence from other systems has demonstrated that calcium channel activity can have a profound effect on quantal size. For example, work at the *C. elegans* NMJ has demonstrated that calcium from intracellular and extracellular sources combines to dictate quantal size and frequency (56). Additionally, spontaneous miniature events with large amplitudes (“maximinis”) have been documented at fast inhibitory synapses of the cerebellum (55, 74). Similar to the NMJ activity documented in our study, these maximinis rely on the ability of ryanodine-sensitive stores to support spontaneous calcium transients large enough to cause multi-vesicular release. It is possible that the architecture of a giant synapse like the Drosophila NMJ – which contains hundreds of active zones clustered into individual boutons and has a low level of spontaneous, multi-vesicular release (75) – makes it exquisitely sensitive to small changes in intracellular calcium from both extracellular and store sources.

*Evoked waveforms*. As is the case for the Drosophila NMJ EPSPs (Fig. 4), the diaphragm NMJ of FHM1 knock-in mice displayed EPP broadening (11). However, the extra discharges we found at the Drosophila NMJ do not seem to be documented for the mammalian NMJ. An instructive parallel may be drawn between our data and cultures of Drosophila giant neurons, in which manipulation of the voltage-gated potassium current generated altered waveforms, including extra and extended discharges (76, 77). It is possible that some aspects of the FHM1 phenotypes may be caused by the perturbation of other voltage-activated currents, and by synapse excitability more generally (78). This possibility is consistent with the fact that mutations in the Na^+^/K^+^ ATPase gene also cause a form of pure FHM (79). Given the effectiveness of the Drosophila system for uncovering complex relationships amongst ion channel activities, in particular potassium currents, the fly may be a good model for studying the cellular bases of disorders such as FHM1 (80–84).

### FHM and non-FHM Migraine: Treatments

Our data suggest that a fly model could uncover molecules that could be targeted to mitigate effects of gain-of-function calcium channel activity associated with migraine. A novel and intriguing finding of our study is factors controlling intracellular calcium store release can be targeted to mitigate FHM1-like hyperexcitability (Figs. 8, 9). Indeed, the RyR channel blocker dantrolene has established uses in the clinic (85, 86). Moreover, significant evidence indicates that blockade of RyR by dantrolene could have neuroprotective applications (87). In the context of FHM1, store operated calcium release would be a novel pathway to consider. Furthermore, lithium (Fig. 9) has been employed in treating migraine, but only in limited cases.

One caveat to our findings is that impairing intracellular calcium release signaling pathways did not reverse all phenotypes associated with SL- and RQ,SL-expressing NMJs. For instance, in the case of increased mEPSP frequency, there is no significant reversal (Table 3). Another caveat is our fly lethality data (Table 4), which suggest that the excitability of single nerve terminals or circuits is not be the only factor to consider. In the case of our fly model, global impairment of factors controlling calcium store release dampened hyperexcitability but enhanced lethality (Table 4). These results point to the fact that gain-of-function Ca_V_2 substitutions may cause multiple, separable, pleiotropic effects. It is possible that the neuronal hyperexcitability phenotypes are somehow protective for fruit fly viability or health – or are a reflection of a protective process that gets blunted when PLCβ and RyR are diminished. Similar considerations could be important in the context of any human migraine treatment.

There is no single, gold-standard pharmaceutical treatment for forms of hemiplegic migraine (88). Several treatments have been employed in clinical settings (89, 90), each with serious drawbacks. Some agents employed to treat hemiplegic migraine include calcium channel blockers like flunarizine (91, 92) and verapamil (Ca_V_1-blocking and potentially Ca_V_2-blocking at higher doses) (93, 94). Blocking of voltage-gated calcium channels would seem to be an intuitive way to counter gain-of-function *CACNA1A* mutations; yet there would be obvious side effects of interfering with Ca_V_2.1 function globally. Other agents reported to be effective in treating hemiplegic migraine are lamotrigine (targeting Na_V_ and Ca_V_2 channels), sodium valproate (several targets including Ca_V_3 channels, resulting in increased inhibitory signaling), and acetazolamide (a pH modulator via carbonic anhydrase inhibition) (see for detailed review (89, 95)). Finally, in cases where hemiplegic migraine attacks are frequent, prophylactic use of triptans has been employed (88, 89). Triptans are a standard treatment for generalized migraine attacks, but since they are vasoconstrictors, there has been some thought that they may not be appropriate for hemiplegic migraine.

Why might a new model be useful, specifically for FHM? FHM is unlike other chronic migraine conditions due its underpinning in central and cortical hyperexcitability and susceptibility to cortical spreading depression (96). In recent years, calcitonin gene-related peptide (CGRP)-based and peripheral approaches have been the focus of generalized migraine treatment. From recent work in mice, there is evidence that CGRP induces migraine-reminiscent photophobia both peripherally and centrally (97). Yet it is uncertain whether CGRP-based therapies would be effective for FHM. For one consideration, CGRP injections do not induce migraine in individuals with FHM in the same manner that it does for other chronic migraineurs sensitive to CGRP levels (98, 99). Recent clinical trials support the use of anti-CGRP receptor antibodies for migraine prophylaxis (100–102), and the Food and Drug Administration (FDA) of the United States has recently approved the anti-CGRP receptor antibody drug Erenumab as a therapeutic (103). Yet the supporting studies did not use individuals with a history of hemiplegic migraine and the antibodies likely act by peripheral mechanisms because they cannot readily cross the blood/brain barrier (104–106). Finally, triptan-based treatments act via reduction in CGRP release and act peripherally where they reverse the effects of CGRP on vasculature (107–109). Given these facts, a new model by which to screen for pharmaco-genetic targets of FHM-causing mutations – such as use of coarse phenotypes of electrophysiological phenotypes in flies – may be valuable.

### Limitations and Future Directions

One strength of Drosophila is the power of genetic manipulation. The genetic toolkit afforded to Drosophila neuroscience makes the NMJ a useful model synapse. One caution regarding the model we generated for this study is that it utilizes over-expression of wild-type or mutant *UAS-cacophony* transgenes. The wild-type version of this transgene recapitulates wild-type *cac* function without generating hyper-excitability phenotypes (22, 28), and we also controlled for potential overexpression phenotypes (Fig. 2). Nevertheless, downstream analyses can be obscured by the need to separate over-expression (hypermorphic) gain-of-function versus mutant (neomorphic) gain-of-function analyses. Other methods, such as CRISPR-based knock-in mutations or expression of a genomic *cac* construct (as employed in (110)) could yield expression levels more similar to endogenous *cac*. Although mutations in the endogenous *cac* locus would be advantageous, we do not expect that these particular limitations detract from our core findings.

We have shown that genetic or pharmacological impairment of an intracellular calcium release signaling pathway suppresses some gain-of-function Ca_V_2 electrophysiological phenotypes. Yet the precise mechanism and sequence of events underlying hyperexcitability suppression we observe are unclear. Potentiation of the baseline activities of the IP_3_R and RyR channels by mutant Ca_V_2 channels is one possibility (111–114). In principle, potentiated RyR or IP_3_R activity could feedback to and further potentiate Ca_V_2 channels. Another possibility is that these gain of function mutations result in chronically increased of intracellular [Ca^2+^] (as in (12)), which could then be reversed indirectly by targeting store pathways. Yet another possibility is that impairment of Ca^2+^ store-release mechanisms somehow dampens Ca_V_2 gating functions – effectively reversing gating gains of function that result from FHM1-causing mutations. Many future directions are possible, utilizing reagents that exist for Drosophila work. A mechanistic refinement could be aided by visual data – for instance by combining inhibition of Ca^2+^ store release along with visual analyses of action potential waveforms via voltage imaging (115) and measurements of Ca_V_2-mediated influx of Ca^2+^ via genetically-encoded indicators tethered to active zone sites (116) – and separately by examining Ca^2+^ dynamics at the stores themselves by using an ER-localizing, genetically encoded calcium sensor developed for Drosophila (117).

The implication of PLCβ activity and intracellular calcium in hyperexcitability is novel within the context of FHM1 mutations, but in hindsight, it also fits with results of prior studies. One recent RNA profiling analysis of the cerebellum of SL knock-in mice revealed an overrepresentation of several signaling components, including PLCβ (118). Moreover, PLCβ and the release of calcium from intracellular stores have been implicated in signaling by CGRP (119–121), whose levels are correlated with generalized migraine (122–124). Beyond work in the Drosophila model, further investigation will be needed to establish whether there is actually a causative link between the action of intracellular calcium stores either in inducing migraine or in precipitating neurological events that precede some forms of migraine, like aura and cortical spreading depression.

## MATERIALS AND METHODS

### Gain-of-function *cacophony* constructs

To generate *UAS-cac-eGFP^SL^* transgenes, we used PCR to alter the serine 161 codon to leucine in the pUAST-based *UAS-cac-eGFP* DNA construct (20, 22). This substitution corresponds to S218L in mammalian CACNA1A. To generate *UAS-cac-eGFP^RQ^* transgenes, we used PCR to change the arginine 135 codon to glutamine. This substitution corresponds to R192Q in mammalian CACNA1A. For the *UAS-cac-eGFP^RQ,SL^* transgene, both mutations were incorporated into the same *UAS-cac-eGFP* construct using PCR to link the overlapping RQ and SL fragments. Transgenic lines were generated by injection of *UAS-cac-eGFP* constructs into a *w^1118^* background (The Best Gene, Chino Hills, CA) and mapped and backcrossed.

### Drosophila Stocks, Genetics, and Husbandry

Animals used for viability counts and electrophysiology were generated by driving neuronal expression of *UAS-cac-eGFP* transgenes with *elaV(C155)-Gal4* (23). Multiple *UAS-cac-eGFP* transgenic lines were initially examined to control for possible differences caused by independent *UAS* genomic insertions: WT: *UAS-cac-eGFP^786c^* (22), *UAS-cac-eGFP^422a^* (22); SL: *UAS-cac-eGFP^S/L(3-2M)^*, *UAS-cac-eGFP^S/L(3-6M)^*, *UAS-cac-eGFP^S/L(3-8M)^*; RQ: *UAS-cac-eGFP^R/Q(1M)^*, *UAS-cac-eGFP^R/Q(2-4M)^*; RQ,SL: *UAS-cac-eGFP^RQ,SL(1M)^*, *UAS-cac-eGFP^RQ,SL(2M)^*.

*w^1118^* (125) was used as a non-transgenic wild-type control. Other Drosophila mutant alleles used were *Df2L(BSC4)* (K. Cook to flybase.org), *Plc21C^p60A^* (126), *Plc21C^MI01911^* (127), *itpr^ug3^* (128), *RyR^E4340K^* (67), *RyR^R4305C^* (67), *RyR^k0943^* (129), *RyR^16^* (130), *RyR^Q3878X^* (67), *RyR^Y4452X^* (67), *Gq^28^* (131), *Gq^221c^* (132), and *cac^HC129^* (35). Mutant Drosophila stocks were obtained either from the Bloomington Drosophila Stock Center (BDSC, Bloomington, Indiana) or directly from the labs that generated them. The *UAS-Plc21C(RNAi)* transformant lines 26557 and 26558 (*Plc21C^GD11359^*) (133) were obtained from the Vienna Drosophila Resource Center (VDRC, Vienna, Austria). A *Gal80^TS^* expression line (39) was employed for a temporal *Gal4* expression experiment. Flies were raised at 25°C (or 29°C for one temperature shift experiment) in humidity- and light-controlled Percival incubators (Geneva Scientific, Fontana, WI), in glass vials on a standard Drosophila food containing water, agar, molasses, yellow cornmeal, and yeast.

### Electrophysiology and Analysis

Wandering third-instar larvae were selected for analysis. Larvae were dissected in a modified HL3 saline with the following components (and concentrations): NaCl (70 mM), KCl (5 mM), MgCl_2_ (10 mM or 6 mM as noted), NaHCO_3_ (10 mM), sucrose (115 mM = 3.9%), trehalose (4.2 mM = 0.16%), HEPES (5.0 mM = 0.12%), and CaCl_2_ (0.5 mM, unless otherwise noted). The central nervous system was removed, except for specific instances noted (Figures 4 and 6). Pharmacological agents tetrodotoxin (TTX, Tocris/R&D Systems), BAPTA-AM (Sigma), Xestospongin C (Tocris/R&D), or lithium chloride (LiCl, Sigma) were added as noted for some experiments. For the experiment using TTX (select agent toxin), all appropriate federal regulations and protocols established for the Select Agent Program established by the Centers for Disease Control and Prevention (CDC) and the US Department of Agriculture (USDA) were followed.

Electrophysiological data were collected using Axopatch 200B or Axoclamp 900A amplifiers (Molecular Devices, Sunnyvale, CA). Sharp electrode (> 10 MΩ) recordings were taken from muscle 6 of abdominal segments 2 and 3, as described previously (30, 31, 134). Prior to muscle V_m_ measurements, the Axoclamp 900A was bridge balanced. For the Axopatch 200B, the amplifier was placed in bridge mode (using I-CLAMP FAST for sharp electrode recordings). Before recording from each muscle, electrode resistance was measured and properly compensated by applying a step input and adjusting series resistance. Muscles with a V_m_ more hyperpolarized than −60 mV and an input resistance of greater than 5 MΩ were deemed suitable for recording (30). Data were digitized using a Digidata 1440A data acquisition system (Molecular Devices) and recorded using the pCLAMP 10 acquisition software (Molecular Devices). Spontaneous activity was recorded, followed by evoked activity. For presynaptic nerve stimulation, a Master-8 pulse stimulator (A.M.P. Instruments, Jerusalem, Israel) and an ISO-Flex isolation unit (A.M.P. Instruments) were utilized to deliver suprathreshold stimuli (1 ms unless otherwise indicated) to the appropriate segmental nerve. For each NMJ, the average amplitude of spontaneous miniature excitatory postsynaptic potential EPSPs (mEPSPs) was quantified by measuring approximately 100-200 individual spontaneous release events per NMJ. The average per-NMJ mEPSP amplitudes were then averaged for each genotype. Evoked EPSP amplitude was calculated for each NMJ as the average of 30 events (1 Hz). Quantal content (QC) was determined in two different ways. At very low extracellular [Ca^2+^], QC was calculated by the method of failures, as *m* = ln[(# trials)/(# failures)], as described elsewhere (42). At higher extracellular [Ca^2+^], QC was calculated by dividing EPSP/mEPSP, as described in the text. For analyses conducted across different calcium concentrations, QC was corrected for non-linear summation (135). For histograms displaying mEPSP amplitude frequencies, the same number of spontaneous events was analyzed for each NMJ (per genotype or experimental condition). This ensured that no individual NMJs were overrepresented or underrepresented in the aggregate analyses.

### Immunostaining and Image Analysis

Third instar larvae were filleted in HL3 saline. Dissected animals were fixed for 3 minutes in Bouin’s fixative (Ricca Chemical Company, Arlington, TX), washed using standard procedures, and incubated in primary antibodies overnight at 4°C. This was followed by additional washes and a two-hour incubation in secondary antibody at room temperature. Staining was performed using the following primary antibodies: mouse anti-GluRIIA (8B4D2) at 1:250 (bouton/cluster counting) or 1:500 (intensity analyses) (Developmental Studies Hybridoma Bank (DSHB), University of Iowa); rabbit anti-Dlg 1:30,000 (136, 137), mouse anti-Brp (nc82) 1:250 (33) (deposited to DSHB by Buchner, E.), rabbit anti-GFP 1:250 (Torrey Pines Biolabs Inc. TP401). The following fluorophore-conjugated antibodies were also used (Jackson ImmunoResearch Laboratories): goat anti-mouse-488 1:1000 (DyLight); and goat anti-rabbit-549 1:2000 (DyLight). Larval preparations were mounted in Vectashield (Vector Laboratories) and imaged at room temperature using Zen software on a Zeiss 700 LSM mounted on an Axio Observer.Z1. An EC Plan-Neofluar 40X Oil DIC Objective (aperture 1.30) or an EC Plan-Apochromat 63x Oil DIC Objective (aperture 1.40) (Carl Zeiss Microscopy) was used.

For analysis of fluorescence intensity and area, experimental and control larval preparations were stained in the same container, mounted on the same slide, imaged using identical acquisition settings, and analyzed using the same procedure and thresholds. Bouton and glutamate receptor cluster numbers were quantified semi-automatically using the ‘Spots’ function in Imaris x64 v7.6.0 (Bitplane, Zurich Switzerland). Any errors in automated counting were corrected by hand to arrive at the final value. GluRIIA and Dlg levels were assessed using ImageJ 1.48s/Java 1.6.0_24 (64-bit) with Fiji plugins. Z-stack images were compressed using the maximum projection function; ROIs were hand drawn to exclude non-synaptic structures; a minimum threshold was set for each channel to eliminate background fluorescence; and the Measure function was used to assess fluorescence intensity and area.

### Western Blotting

10 adult fly heads/sample were prepared in sample buffer using standard methods. SDS-PAGE was performed using the Novex NuPAGE SDS-PAGE system with 4%-12% Bis-Tris gels run at 125 V for 10 minutes and 150 V for 2.5 hours. Transfer to PVDF membrane (Bio-Rad, Hercules, CA) was performed using a Trans-Blot-SDSemi-Dry Transfer Cell (Bio-Rad, Hercules, CA). Blocking was performed in 5% BSA for GFP blots or 5% milk for actin blots in 1X PBS with 0.1% Tween 20. Primary antibodies were obtained from the DSHB, mouse anti-actin (JLA20) 1:1000, or from Torrey Pines Biolabs, rabbit anti-GFP 1:2000. Horseradish peroxidase-conjugated goat anti-mouse secondary antibody (Jackson ImmunoResearch Laboratories, Inc., West Grove, PA) was used at 1:5000 for actin blots. Horseradish peroxidase-conjugated goat anti-rabbit secondary antibody (Jackson ImmunoResearch Laboratories, Inc., West Grove, PA) was used at 1:5000 for GFP blots. All antibodies were diluted in blocking buffer. Blots were developed with Super-Signal West Pico Chemiluminescent Substrate (Thermo Scientific, Waltham, MA) and imagedwith Amersham Hyperfilm ECL film (GE Healthcare Limited, Buckinghamshire, UK). Band intensity was quantified using ImageJ.

### Statistical Analyses and Data Plots

Most electrophysiological comparisons were made across multiple data sets. As appropriate, statistical significance was either assessed by one-way ANOVA with Tukey’s post-hoc analysis for multiple comparisons (assumes Gaussian distribution), or a non-parametric Kruskal-Wallis ANOVA with Dunn’s post-hoc analysis for multiple comparisons (does not assume Gaussian distribution). Other statistical tests utilized included Fisher’s exact tests for viability counts and for counts of gigantic mEPSP events; Log-rank tests for survivability curves; linear regression analyses for calcium cooperativity; and Student’s T-Tests for direct comparisons between one control group and one experimental group. *p* values of * *p* < 0.05, ** *p* < 0.01, *** *p* < 0.001, and **** *p* < 0.0001 were considered significant. The values reported or plotted on regular bar graphs are mean ± SEM. The values reported and plotted on box-and-whisker graphs are: box (25^th^ – 75^th^ percentiles), whiskers (1^st^ – 99^th^ percentiles), line (median), + symbol (average), and individual raw data points plotted outside the 1^st^ and 99^th^ percentiles. Statistical analyses were performed in GraphPad Prism (GraphPad Software).

## ACKNOWLEDGEMENTS

We thank Martin Müller, Chun-Fang Wu and members of the Frank lab for helpful comments on earlier versions of this manuscript, and the laboratories of Tina Tootle and Fang Lin for helpful discussions. We thank the Bloomington Drosophila Stock Center and the Vienna Drosophila Resource Center for several fly stocks detailed in the Materials and Methods section.

## REFERENCES

1. Kullmann DM. Neurological channelopathies. Annu Rev Neurosci. 2010;33:151–72.

2. Russell JF, Fu YH, Ptacek LJ. Episodic neurologic disorders: syndromes, genes, and mechanisms. Annu Rev Neurosci. 2013;36:25–50.

3. Ryan DP, Ptacek LJ. Episodic neurological channelopathies. Neuron. 2010;68(2):282–92.

4. Ophoff RA, Terwindt GM, Vergouwe MN, van Eijk R, Oefner PJ, Hoffman SM, et al. Familial hemiplegic migraine and episodic ataxia type-2 are caused by mutations in the Ca2+ channel gene CACNL1A4. Cell. 1996;87(3):543–52.

5. Kors EE, Terwindt GM, Vermeulen FL, Fitzsimons RB, Jardine PE, Heywood P, et al. Delayed cerebral edema and fatal coma after minor head trauma: role of the CACNA1A calcium channel subunit gene and relationship with familial hemiplegic migraine. Ann Neurol. 2001;49(6):753–60.

6. Pietrobon D. Calcium channels and migraine. Biochimica et biophysica acta. 2013;1828(7):1655–65.

7. Eikermann-Haerter K, Dilekoz E, Kudo C, Savitz SI, Waeber C, Baum MJ, et al. Genetic and hormonal factors modulate spreading depression and transient hemiparesis in mouse models of familial hemiplegic migraine type 1. The Journal of clinical investigation. 2009;119(1):99–109.

8. van den Maagdenberg AM, Pietrobon D, Pizzorusso T, Kaja S, Broos LA, Cesetti T, et al. A Cacna1a knockin migraine mouse model with increased susceptibility to cortical spreading depression. Neuron. 2004;41(5):701–10.

9. van den Maagdenberg AM, Pizzorusso T, Kaja S, Terpolilli N, Shapovalova M, Hoebeek FE, et al. High cortical spreading depression susceptibility and migraine-associated symptoms in Ca(v)2.1 S218L mice. Ann Neurol. 2010;67(1):85–98.

10. Kaja S, van de Ven RC, Broos LA, Veldman H, van Dijk JG, Verschuuren JJ, et al. Gene dosage-dependent transmitter release changes at neuromuscular synapses of CACNA1A R192Q knockin mice are non-progressive and do not lead to morphological changes or muscle weakness. Neuroscience. 2005;135(1):81–95.

11. Kaja S, Van de Ven RC, Broos LA, Frants RR, Ferrari MD, Van den Maagdenberg AM, et al. Severe and progressive neurotransmitter release aberrations in familial hemiplegic migraine type 1 Cacna1a S218L knock-in mice. Journal of neurophysiology. 2010;104(3):1445–55.

12. Di Guilmi MN, Wang T, Inchauspe CG, Forsythe ID, Ferrari MD, van den Maagdenberg AM, et al. Synaptic Gain-of-Function Effects of Mutant Cav2.1 Channels in a Mouse Model of Familial Hemiplegic Migraine Are Due to Increased Basal [Ca2+]i. J Neurosci. 2014;34(21):7047–58.

13. Inchauspe CG, Urbano FJ, Di Guilmi MN, Ferrari MD, van den Maagdenberg AM, Forsythe ID, et al. Presynaptic CaV2.1 calcium channels carrying familial hemiplegic migraine mutation R192Q allow faster recovery from synaptic depression in mouse calyx of Held. Journal of neurophysiology. 2012;108(11):2967–76.

14. Inchauspe CG, Urbano FJ, Di Guilmi MN, Forsythe ID, Ferrari MD, van den Maagdenberg AM, et al. Gain of function in FHM-1 Cav2.1 knock-in mice is related to the shape of the action potential. Journal of neurophysiology. 2010;104(1):291–9.

15. Hullugundi SK, Ansuini A, Ferrari MD, van den Maagdenberg AM, Nistri A. A hyperexcitability phenotype in mouse trigeminal sensory neurons expressing the R192Q Cacna1a missense mutation of familial hemiplegic migraine type-1. Neuroscience. 2014;266:244–54.

16. Park J, Moon H, Akerman S, Holland PR, Lasalandra MP, Andreou AP, et al. Differential trigeminovascular nociceptive responses in the thalamus in the familial hemiplegic migraine 1 knock-in mouse: a Fos protein study. Neurobiology of disease. 2014;64:1–7.

17. Pietrobon D, Moskowitz MA. Pathophysiology of migraine. Annu Rev Physiol. 2013;75:365–91.

18. Vecchia D, Tottene A, van den Maagdenberg AM, Pietrobon D. Abnormal cortical synaptic transmission in CaV2.1 knockin mice with the S218L missense mutation which causes a severe familial hemiplegic migraine syndrome in humans. Frontiers in cellular neuroscience. 2015;9:8.

19. Vecchia D, Tottene A, van den Maagdenberg AM, Pietrobon D. Mechanism underlying unaltered cortical inhibitory synaptic transmission in contrast with enhanced excitatory transmission in CaV2.1 knockin migraine mice. Neurobiology of disease. 2014;69:225–34.

20. Inagaki A, Frank CA, Usachev YM, Benveniste M, Lee A. Pharmacological Correction of Gating Defects in the Voltage-Gated Cav2.1 Ca(2+) Channel due to a Familial Hemiplegic Migraine Mutation. Neuron. 2014;81(1):91–102.

21. Eikermann-Haerter K, Arbel-Ornath M, Yalcin N, Yu ES, Kuchibhotla KV, Yuzawa I, et al. Abnormal synaptic Ca(2+) homeostasis and morphology in cortical neurons of familial hemiplegic migraine type 1 mutant mice. Ann Neurol. 2015;78(2):193–210.

22. Kawasaki F, Zou B, Xu X, Ordway RW. Active zone localization of presynaptic calcium channels encoded by the cacophony locus of Drosophila. J Neurosci. 2004;24(1):282–5.

23. Lin DM, Goodman CS. Ectopic and increased expression of Fasciclin II alters motoneuron growth cone guidance. Neuron. 1994;13(3):507–23.

24. Brand AH, Perrimon N. Targeted gene expression as a means of altering cell fates and generating dominant phenotypes. Development. 1993;118(2):401–15.

25. Breen TR, Lucchesi JC. Analysis of the dosage compensation of a specific transcript in Drosophila melanogaster. Genetics. 1986;112(3):483–91.

26. Cline TW, Meyer BJ. Vive la difference: males vs females in flies vs worms. Annual review of genetics. 1996;30:637–702.

27. Lucchesi JC, Kuroda MI. Dosage compensation in Drosophila. Cold Spring Harbor perspectives in biology. 2015;7(5).

28. Kawasaki F, Collins SC, Ordway RW. Synaptic calcium-channel function in Drosophila: analysis and transformation rescue of temperature-sensitive paralytic and lethal mutations of cacophony. J Neurosci. 2002;22(14):5856–64.

29. Fouquet W, Owald D, Wichmann C, Mertel S, Depner H, Dyba M, et al. Maturation of active zone assembly by Drosophila Bruchpilot. J Cell Biol. 2009;186(1):129–45.

30. Frank CA, Kennedy MJ, Goold CP, Marek KW, Davis GW. Mechanisms underlying the rapid induction and sustained expression of synaptic homeostasis. Neuron. 2006;52(4):663–77.

31. Frank CA, Pielage J, Davis GW. A presynaptic homeostatic signaling system composed of the Eph receptor, ephexin, Cdc42, and CaV2.1 calcium channels. Neuron. 2009;61(4):556–69.

32. Gaviño MA, Ford KJ, Archila S, Davis GW. Homeostatic synaptic depression is achieved through a regulated decrease in presynaptic calcium channel abundance. eLife. 2015;4.

33. Wagh DA, Rasse TM, Asan E, Hofbauer A, Schwenkert I, Durrbeck H, et al. Bruchpilot, a protein with homology to ELKS/CAST, is required for structural integrity and function of synaptic active zones in Drosophila. Neuron. 2006;49(6):833–44.

34. DiAntonio A, Petersen SA, Heckmann M, Goodman CS. Glutamate receptor expression regulates quantal size and quantal content at the Drosophila neuromuscular junction. J Neurosci. 1999;19(8):3023–32.

35. Kulkarni SJ, Hall JC. Behavioral and cytogenetic analysis of the cacophony courtship song mutant and interacting genetic variants in Drosophila melanogaster. Genetics. 1987;115(3):461–75.

36. Feng Y, Ueda A, Wu CF. A modified minimal hemolymph-like solution, HL3.1, for physiological recordings at the neuromuscular junctions of normal and mutant Drosophila larvae. Journal of neurogenetics. 2004;18(2):377–402.

37. Tottene A, Conti R, Fabbro A, Vecchia D, Shapovalova M, Santello M, et al. Enhanced excitatory transmission at cortical synapses as the basis for facilitated spreading depression in Ca(v)2.1 knockin migraine mice. Neuron. 2009;61(5):762–73.

38. Adams PJ, Rungta RL, Garcia E, van den Maagdenberg AM, MacVicar BA, Snutch TP. Contribution of calcium-dependent facilitation to synaptic plasticity revealed by migraine mutations in the P/Q-type calcium channel. Proc Natl Acad Sci U S A. 2010;107(43):18694–9.

39. McGuire SE, Le PT, Osborn AJ, Matsumoto K, Davis RL. Spatiotemporal rescue of memory dysfunction in Drosophila. Science. 2003;302(5651):1765–8.

40. Jan LY, Jan YN. Properties of the larval neuromuscular junction in Drosophila melanogaster. J Physiol. 1976;262(1):189–214.

41. Martin AR. Quantal Nature of Synaptic Transmission. Physiol Rev. 1966;46:51–66.

42. Del Castillo J, Katz B. Quantal components of the end-plate potential. J Physiol. 1954;124(3):560–73.

43. Frank CA. Homeostatic plasticity at the Drosophila neuromuscular junction. Neuropharmacology. 2014;78:63–74.

44. Frank CA. How voltage-gated calcium channels gate forms of homeostatic synaptic plasticity. Frontiers in cellular neuroscience. 2014;8:40.

45. Brusich DJ, Spring AM, Frank CA. A single-cross, RNA interference-based genetic tool for examining the long-term maintenance of homeostatic plasticity. Frontiers in cellular neuroscience. 2015;9:107.

46. Shortridge RD, Yoon J, Lending CR, Bloomquist BT, Perdew MH, Pak WL. A Drosophila phospholipase C gene that is expressed in the central nervous system. J Biol Chem. 1991;266(19):12474–80.

47. Tedford HW, Zamponi GW. Direct G protein modulation of Cav2 calcium channels. Pharmacol Rev. 2006;58(4):837–62.

48. Cremona O, De Camilli P. Phosphoinositides in membrane traffic at the synapse. Journal of cell science. 2001;114(Pt 6):1041–52.

49. Goni FM, Alonso A. Structure and functional properties of diacylglycerols in membranes. Progress in lipid research. 1999;38(1):1–48.

50. Huang FD, Woodruff E, Mohrmann R, Broadie K. Rolling blackout is required for synaptic vesicle exocytosis. J Neurosci. 2006;26(9):2369–79.

51. Peters C, Bayer MJ, Buhler S, Andersen JS, Mann M, Mayer A. Trans-complex formation by proteolipid channels in the terminal phase of membrane fusion. Nature. 2001;409(6820):581–8.

52. Rohrbough J, Broadie K. Lipid regulation of the synaptic vesicle cycle. Nat Rev Neurosci. 2005;6(2):139–50.

53. Wu L, Bauer CS, Zhen XG, Xie C, Yang J. Dual regulation of voltage-gated calcium channels by PtdIns(4,5)P2. Nature. 2002;419(6910):947–52.

54. Dahdal D, Reeves DC, Ruben M, Akabas MH, Blau J. Drosophila pacemaker neurons require g protein signaling and GABAergic inputs to generate twenty-four hour behavioral rhythms. Neuron. 2010;68(5):964–77.

55. Llano I, Gonzalez J, Caputo C, Lai FA, Blayney LM, Tan YP, et al. Presynaptic calcium stores underlie large-amplitude miniature IPSCs and spontaneous calcium transients. Nat Neurosci. 2000;3(12):1256–65.

56. Liu Q, Chen B, Yankova M, Morest DK, Maryon E, Hand AR, et al. Presynaptic ryanodine receptors are required for normal quantal size at the Caenorhabditis elegans neuromuscular junction. J Neurosci. 2005;25(29):6745–54.

57. Collin T, Marty A, Llano I. Presynaptic calcium stores and synaptic transmission. Curr Opin Neurobiol. 2005;15(3):275–81.

58. Emptage NJ, Reid CA, Fine A. Calcium stores in hippocampal synaptic boutons mediate short-term plasticity, store-operated Ca2+ entry, and spontaneous transmitter release. Neuron. 2001;29(1):197–208.

59. Simkus CR, Stricker C. The contribution of intracellular calcium stores to mEPSCs recorded in layer II neurones of rat barrel cortex. J Physiol. 2002;545(Pt 2):521–35.

60. Wong CO, Chen K, Lin YQ, Chao Y, Duraine L, Lu Z, et al. A TRPV channel in Drosophila motor neurons regulates presynaptic resting Ca2+ levels, synapse growth, and synaptic transmission. Neuron. 2014;84(4):764–77.

61. Joshi R, Venkatesh K, Srinivas R, Nair S, Hasan G. Genetic dissection of itpr gene function reveals a vital requirement in aminergic cells of Drosophila larvae. Genetics. 2004;166(1):225–36.

62. Conti R, Tan YP, Llano I. Action potential-evoked and ryanodine-sensitive spontaneous Ca2+ transients at the presynaptic terminal of a developing CNS inhibitory synapse. J Neurosci. 2004;24(31):6946–57.

63. Dunn TW, Syed NI. Ryanodine receptor-transmitter release site coupling increases quantal size in a synapse-specific manner. The European journal of neuroscience. 2006;24(6):1591–605.

64. Gordon GR, Bains JS. Noradrenaline triggers multivesicular release at glutamatergic synapses in the hypothalamus. J Neurosci. 2005;25(49):11385–95.

65. Sharma G, Vijayaraghavan S. Modulation of presynaptic store calcium induces release of glutamate and postsynaptic firing. Neuron. 2003;38(6):929–39.

66. Dunn TW, McCamphill PK, Syed NI. Activity-induced large amplitude postsynaptic mPSPs at soma-soma synapses between Lymnaea neurons. Synapse. 2009;63(2):117–25.

67. Dockendorff TC, Robertson SE, Faulkner DL, Jongens TA. Genetic characterization of the 44D-45B region of the Drosophila melanogaster genome based on an F2 lethal screen. Molecular & general genetics: MGG. 2000;263(1):137–43.

68. Hallcher LM, Sherman WR. The effects of lithium ion and other agents on the activity of myo-inositol-1-phosphatase from bovine brain. J Biol Chem. 1980;255(22):10896–901.

69. Lenox RH, Wang L. Molecular basis of lithium action: integration of lithium-responsive signaling and gene expression networks. Mol Psychiatry. 2003;8(2):135–44.

70. Gafni J, Munsch JA, Lam TH, Catlin MC, Costa LG, Molinski TF, et al. Xestospongins: potent membrane permeable blockers of the inositol 1,4,5-trisphosphate receptor. Neuron. 1997;19(3):723–33.

71. Wilcox RA, Primrose WU, Nahorski SR, Challiss RA. New developments in the molecular pharmacology of the myo-inositol 1,4,5-trisphosphate receptor. Trends Pharmacol Sci. 1998;19(11):467–75.

72. Goswami SP, Bucurenciu I, Jonas P. Miniature IPSCs in hippocampal granule cells are triggered by voltage-gated Ca2+ channels via microdomain coupling. J Neurosci. 2012;32(41):14294–304.

73. Ermolyuk YS, Alder FG, Surges R, Pavlov IY, Timofeeva Y, Kullmann DM, et al. Differential triggering of spontaneous glutamate release by P/Q-, N- and R-type Ca2+ channels. Nat Neurosci. 2013;16(12):1754–63.

74. Xu-Friedman MA, Regehr WG. Maximinis. Nat Neurosci. 2000;3(12):1229–30.

75. Melom JE, Akbergenova Y, Gavornik JP, Littleton JT. Spontaneous and evoked release are independently regulated at individual active zones. J Neurosci. 2013;33(44):17253–63.

76. Yao WD, Wu CF. Auxiliary Hyperkinetic beta subunit of K+ channels: regulation of firing properties and K+ currents in Drosophila neurons. J Neurophysiol. 1999;81(5):2472–84.

77. Berke BA, Lee J, Peng IF, Wu CF. Sub-cellular Ca2+ dynamics affected by voltage- and Ca2+- gated K+ channels: Regulation of the soma-growth cone disparity and the quiescent state in Drosophila neurons. Neuroscience. 2006;142(3):629–44.

78. Gao Z, Todorov B, Barrett CF, van Dorp S, Ferrari MD, van den Maagdenberg AM, et al. Cerebellar ataxia by enhanced Ca(V)2.1 currents is alleviated by Ca2+-dependent K+-channel activators in Cacna1a(S218L) mutant mice. J Neurosci. 2012;32(44):15533–46.

79. Vanmolkot KR, Kors EE, Turk U, Turkdogan D, Keyser A, Broos LA, et al. Two de novo mutations in the Na, K-ATPase gene ATP1A2 associated with pure familial hemiplegic migraine. European journal of human genetics: EJHG. 2006;14(5):555–60.

80. Bergquist S, Dickman DK, Davis GW. A hierarchy of cell intrinsic and target-derived homeostatic signaling. Neuron. 2010;66(2):220–34.

81. Lee J, Ueda A, Wu CF. Pre- and post-synaptic mechanisms of synaptic strength homeostasis revealed by slowpoke and shaker K+ channel mutations in Drosophila. Neuroscience. 2008;154(4):1283–96.

82. Lee J, Ueda A, Wu CF. Distinct roles of Drosophila cacophony and Dmca1D Ca(2+) channels in synaptic homeostasis: genetic interactions with slowpoke Ca(2+) -activated BK channels in presynaptic excitability and postsynaptic response. Developmental neurobiology. 2014;74(1):1–15.

83. Peng IF, Wu CF. Drosophila cacophony channels: a major mediator of neuronal Ca2+ currents and a trigger for K+ channel homeostatic regulation. J Neurosci. 2007;27(5):1072–81.

84. Parrish JZ, Kim CC, Tang L, Bergquist S, Wang T, Derisi JL, et al. Kruppel mediates the selective rebalancing of ion channel expression. Neuron. 2014;82(3):537–44.

85. Coronado R, Morrissette J, Sukhareva M, Vaughan DM. Structure and function of ryanodine receptors. The American journal of physiology. 1994;266(6 Pt 1):C1485–504.

86. Krause T, Gerbershagen MU, Fiege M, Weisshorn R, Wappler F. Dantrolene--a review of its pharmacology, therapeutic use and new developments. Anaesthesia. 2004;59(4):364–73.

87. Muehlschlegel S, Sims JR. Dantrolene: mechanisms of neuroprotection and possible clinical applications in the neurointensive care unit. Neurocrit Care. 2009;10(1):103–15.

88. Jen JC. Familial Hemiplegic Migraine. In: Adam MP, Ardinger HH, Pagon RA, Wallace SE, Bean LJH, Stephens K, et al., editors. GeneReviews((R)). Seattle (WA)1993.

89. Pelzer N, Stam AH, Haan J, Ferrari MD, Terwindt GM. Familial and sporadic hemiplegic migraine: diagnosis and treatment. Curr Treat Options Neurol. 2013;15(1):13–27.

90. Pelzer N, Haan J, Stam AH, Vijfhuizen LS, Koelewijn SC, Smagge A, et al. Clinical spectrum of hemiplegic migraine and chances of finding a pathogenic mutation. Neurology. 2018;90(7):e575–e82.

91. Ye Q, Yan LY, Xue LJ, Wang Q, Zhou ZK, Xiao H, et al. Flunarizine blocks voltage-gated Na(+) and Ca(2+) currents in cultured rat cortical neurons: A possible locus of action in the prevention of migraine. Neurosci Lett. 2011;487(3):394–9.

92. Peer Mohamed B, Goadsby PJ, Prabhakar P. Safety and efficacy of flunarizine in childhood migraine: 11 years’ experience, with emphasis on its effect in hemiplegic migraine. Dev Med Child Neurol. 2012;54(3):274–7.

93. Dobrev D, Milde AS, Andreas K, Ravens U. The effects of verapamil and diltiazem on N-, P- and Q-type calcium channels mediating dopamine release in rat striatum. Br J Pharmacol. 1999;127(2):576–82.

94. Yu W, Horowitz SH. Treatment of sporadic hemiplegic migraine with calcium-channel blocker verapamil. Neurology. 2003;60(1):120–1.

95. Pelzer N, Stam AH, Carpay JA, Vries BD, van den Maagdenberg AM, Ferrari MD, et al. Familial hemiplegic migraine treated by sodium valproate and lamotrigine. Cephalalgia: an international journal of headache. 2014;34(9):708–11.

96. Pietrobon D. Familial hemiplegic migraine. Neurotherapeutics. 2007;4(2):274–84.

97. Mason BN, Kaiser EA, Kuburas A, Loomis MM, Latham JA, Garcia-Martinez LF, et al. Induction of Migraine-Like Photophobic Behavior in Mice by Both Peripheral and Central CGRP Mechanisms. J Neurosci. 2017;37(1):204–16.

98. Hansen JM, Hauge AW, Olesen J, Ashina M. Calcitonin gene-related peptide triggers migraine-like attacks in patients with migraine with aura. Cephalalgia: an international journal of headache. 2010;30(10):1179–86.

99. Hansen JM, Thomsen LL, Olesen J, Ashina M. Calcitonin gene-related peptide does not cause the familial hemiplegic migraine phenotype. Neurology. 2008;71(11):841–7.

100. Goadsby PJ, Reuter U, Hallstrom Y, Broessner G, Bonner JH, Zhang F, et al. A Controlled Trial of Erenumab for Episodic Migraine. The New England journal of medicine. 2017;377(22):2123–32.

101. Tepper S, Ashina M, Reuter U, Brandes JL, Dolezil D, Silberstein S, et al. Safety and efficacy of erenumab for preventive treatment of chronic migraine: a randomised, double-blind, placebo-controlled phase 2 trial. Lancet neurology. 2017;16(6):425–34.

102. Dodick DW, Ashina M, Brandes JL, Kudrow D, Lanteri-Minet M, Osipova V, et al. ARISE: A Phase 3 randomized trial of erenumab for episodic migraine. Cephalalgia: an international journal of headache. 2018;38(6):1026–37.

103. Traynor K. FDA approves licensing of erenumab-aooe to prevent migraine. Am J Health Syst Pharm. 2018;75(13):929–30.

104. Dodick DW, Goadsby PJ, Spierings EL, Scherer JC, Sweeney SP, Grayzel DS. Safety and efficacy of LY2951742, a monoclonal antibody to calcitonin gene-related peptide, for the prevention of migraine: a phase 2, randomised, double-blind, placebo-controlled study. Lancet neurology. 2014;13(9):885–92.

105. Tfelt-Hansen P. Site of effect of LY2951742 for migraine prophylaxis. Lancet neurology. 2015;14(1):31–2.

106. Iyengar S, Ossipov MH, Johnson KW. The role of calcitonin gene-related peptide in peripheral and central pain mechanisms including migraine. Pain. 2017;158(4):543–59.

107. Asghar MS, Hansen AE, Kapijimpanga T, van der Geest RJ, van der Koning P, Larsson HB, et al. Dilation by CGRP of middle meningeal artery and reversal by sumatriptan in normal volunteers. Neurology. 2010;75(17):1520–6.

108. Amrutkar DV, Ploug KB, Hay-Schmidt A, Porreca F, Olesen J, Jansen-Olesen I. mRNA expression of 5-hydroxytryptamine 1B, 1D, and 1F receptors and their role in controlling the release of calcitonin gene-related peptide in the rat trigeminovascular system. Pain. 2012;153(4):830–8.

109. Benemei S, Cortese F, Labastida-Ramirez A, Marchese F, Pellesi L, Romoli M, et al. Triptans and CGRP blockade - impact on the cranial vasculature. J Headache Pain. 2017;18(1):103.

110. Luo X, Rosenfeld JA, Yamamoto S, Harel T, Zuo Z, Hall M, et al. Clinically severe CACNA1A alleles affect synaptic function and neurodegeneration differentially. PLoS genetics. 2017;13(7):e1006905.

111. Li P, Chen SR. Molecular basis of Ca(2)+ activation of the mouse cardiac Ca(2)+ release channel (ryanodine receptor). The Journal of general physiology. 2001;118(1):33–44.

112. Rahman T. Dynamic clustering of IP3 receptors by IP3. Biochemical Society transactions. 2012;40(2):325–30.

113. Zahradnik I, Gyorke S, Zahradnikova A. Calcium activation of ryanodine receptor channels--reconciling RyR gating models with tetrameric channel structure. The Journal of general physiology. 2005;126(5):515–27.

114. Laver DR. Ca2+ stores regulate ryanodine receptor Ca2+ release channels via luminal and cytosolic Ca2+ sites. Clinical and experimental pharmacology & physiology. 2007;34(9):889–96.

115. Ford KJ, Davis GW. Archaerhodopsin voltage imaging: synaptic calcium and BK channels stabilize action potential repolarization at the Drosophila neuromuscular junction. J Neurosci. 2014;34(44):14517–25.

116. Kiragasi B, Wondolowski J, Li Y, Dickman DK. A Presynaptic Glutamate Receptor Subunit Confers Robustness to Neurotransmission and Homeostatic Potentiation. Cell reports. 2017;19(13):2694–706.

117. Navas-Navarro P, Rojo-Ruiz J, Rodriguez-Prados M, Ganfornina MD, Looger LL, Alonso MT, et al. GFP-Aequorin Protein Sensor for Ex Vivo and In Vivo Imaging of Ca(2+) Dynamics in High-Ca(2+) Organelles. Cell Chem Biol. 2016;23(6):738–45.

118. de Vries B, Eising E, Broos LA, Koelewijn SC, Todorov B, Frants RR, et al. RNA expression profiling in brains of familial hemiplegic migraine type 1 knock-in mice. Cephalalgia: an international journal of headache. 2014;34(3):174–82.

119. Drissi H, Lasmoles F, Le Mellay V, Marie PJ, Lieberherr M. Activation of phospholipase C-beta1 via Galphaq/11 during calcium mobilization by calcitonin gene-related peptide. J Biol Chem. 1998;273(32):20168–74.

120. Morara S, Wang LP, Filippov V, Dickerson IM, Grohovaz F, Provini L, et al. Calcitonin gene-related peptide (CGRP) triggers Ca2+ responses in cultured astrocytes and in Bergmann glial cells from cerebellar slices. Eur J Neurosci. 2008;28(11):2213–20.

121. Drissi H, Lieberherr M, Hott M, Marie PJ, Lasmoles F. Calcitonin gene-related peptide (CGRP) increases intracellular free Ca2+ concentrations but not cyclic AMP formation in CGRP receptor-positive osteosarcoma cells (OHS-4). Cytokine. 1999;11(3):200–7.

122. Goadsby PJ, Edvinsson L, Ekman R. Release of vasoactive peptides in the extracerebral circulation of humans and the cat during activation of the trigeminovascular system. Ann Neurol. 1988;23(2):193–6.

123. Goadsby PJ, Edvinsson L, Ekman R. Vasoactive peptide release in the extracerebral circulation of humans during migraine headache. Ann Neurol. 1990;28(2):183–7.

124. Kaiser EA, Russo AF. CGRP and migraine: could PACAP play a role too? Neuropeptides. 2013;47(6):451–61.

125. Hazelrigg T, Levis R, Rubin GM. Transformation of white locus DNA in drosophila: dosage compensation, zeste interaction, and position effects. Cell. 1984;36(2):469–81.

126. Weinkove D, Neufeld TP, Twardzik T, Waterfield MD, Leevers SJ. Regulation of imaginal disc cell size, cell number and organ size by Drosophila class I(A) phosphoinositide 3-kinase and its adaptor. Curr Biol. 1999;9(18):1019–29.

127. Venken KJ, Schulze KL, Haelterman NA, Pan H, He Y, Evans-Holm M, et al. MiMIC: a highly versatile transposon insertion resource for engineering Drosophila melanogaster genes. Nature methods. 2011;8(9):737–43.

128. Deshpande M, Venkatesh K, Rodrigues V, Hasan G. The inositol 1,4,5-trisphosphate receptor is required for maintenance of olfactory adaptation in Drosophila antennae. J Neurobiol. 2000;43(3):282–8.

129. Spradling AC, Stern D, Beaton A, Rhem EJ, Laverty T, Mozden N, et al. The Berkeley Drosophila Genome Project gene disruption project: Single P-element insertions mutating 25% of vital Drosophila genes. Genetics. 1999;153(1):135–77.

130. Sullivan KM, Scott K, Zuker CS, Rubin GM. The ryanodine receptor is essential for larval development in Drosophila melanogaster. Proc Natl Acad Sci U S A. 2000;97(11):5942–7.

131. Yao CA, Carlson JR. Role of G-proteins in odor-sensing and CO2-sensing neurons in Drosophila. J Neurosci. 2010;30(13):4562–72.

132. Banerjee S, Joshi R, Venkiteswaran G, Agrawal N, Srikanth S, Alam F, et al. Compensation of inositol 1,4,5-trisphosphate receptor function by altering sarco-endoplasmic reticulum calcium ATPase activity in the Drosophila flight circuit. J Neurosci. 2006;26(32):8278–88.

133. Dietzl G, Chen D, Schnorrer F, Su KC, Barinova Y, Fellner M, et al. A genome-wide transgenic RNAi library for conditional gene inactivation in Drosophila. Nature. 2007;448(7150):151–6.

134. Davis GW, DiAntonio A, Petersen SA, Goodman CS. Postsynaptic PKA controls quantal size and reveals a retrograde signal that regulates presynaptic transmitter release in Drosophila. Neuron. 1998;20(2):305–15.

135. Martin AR. A further study of the statistical composition on the end-plate potential. J Physiol. 1955;130(1):114–22.

136. Budnik V, Koh YH, Guan B, Hartmann B, Hough C, Woods D, et al. Regulation of synapse structure and function by the Drosophila tumor suppressor gene dlg. Neuron. 1996;17(4):627–40.

137. Pielage J, Cheng L, Fetter RD, Carlton PM, Sedat JW, Davis GW. A presynaptic giant ankyrin stabilizes the NMJ through regulation of presynaptic microtubules and transsynaptic cell adhesion. Neuron. 2008;58(2):195–209.

